# Metabolic Reprogramming of Human Macrophages Drives the Formation of Hybrid M1/M2 Pro-Regenerative Extracellular Vesicles

**DOI:** 10.64898/2026.01.16.699890

**Authors:** Cansu Gorgun, Paula Klavina, Carolina S. Martins, Cloe Payet, Brenton L. Cavanagh, Marianne Pultar, Matthias Hackl, Annie M. Curtis, David A. Hoey

## Abstract

The coordinated activity of macrophages is essential for bone repair, with pro-inflammatory M1 macrophages driving early responses and anti-inflammatory M2 macrophages supporting later tissue remodeling. While both phenotypes are required, prolonged persistence of either subtype can impair healing, underscoring the correct transition between the two states. Macrophage polarization is closely linked to cellular metabolism, and human macrophages display distinct metabolic profiles. Macrophage-derived extracellular vesicles (EVs) carry bioactive cargo and reflect parental polarization, influencing recipient cell function. This raises critical questions about how metabolic regulation influences human macrophage function, their EVs and their effect on angiogenesis and osteogenesis. This study investigates EVs derived from polarized primary human macrophages and from macrophages exposed to DASA-58, a small molecule which activates the metabolic enzyme pyruvate kinase M2 (PKM2). Alterations in macrophage metabolism modifies the molecular cargo of their EVs, including microRNAs (miRNAs), to modulate regenerative activity. These findings demonstrate that human macrophage-derived EVs exert metabolically dependent effects on angiogenesis and osteogenesis, and that metabolic modulation enables the generation of EVs with hybrid pro-regenerative properties intermediate between M1 and M2. This establishes metabolic reprogramming within human macrophages using small molecules as a strategy to engineer novel phenotypes and EVs for bone repair.

## 1. Introduction

Macrophages, which are key cells of innate immunity are central to tissue repair, whereby they exert plasticity in modulating their phenotype and the release of regenerative factors to temporally orchestrate effective tissue rebuild [1]. Following injury, macrophages first adopt an M1 pro-inflammatory profile, contributing to early inflammatory responses and debris clearance, before transitioning to an anti-inflammatory M2 phenotype that supports later stages of healing. In the context of bone repair, this transition is closely linked to angiogenesis, the formation of new blood vessels that supply the injury site with nutrients and immune cells to support osteogenesis. While M1 macrophages play an important role in initial repair, including pro-angiogenic signaling, their sustained activation can drive chronic inflammation. In contrast, M2 macrophages facilitate the resolution of inflammation and promote tissue remodeling, partly through the secretion of factors that enhance vascularization and support bone repair [2]. Dysregulation of this M1/M2 transition compromises healing. Depletion of M1 macrophages during bone repair delays endochondral ossification, highlighting the requirement of M1 macrophages, while loss of M2 macrophages reduces angiogenesis and impairs ossification [3, 4]. While the contributions of macrophages to tissue repair have been explored, published findings present conflicting evidence concerning the specific benefits of M1 or M2 phenotypes on regenerative processes, particularly in the human context [5–8].

Although the traditional M1/M2 classification has been useful, it oversimplifies the complexity of macrophage biology. When the M1/M2 balance is disrupted, such as what occurs in aging, osteoporosis and diabetes mellitus macrophages remain in the M1 phase and fail to transition to the M2 phenotype delaying healing [9–11]. Interestingly, recent evidence has demonstrated that macrophage phenotype and function is closely intertwined with their metabolic state, where macrophage metabolic dysregulation directs the macrophage phenotype [12]. Glycolysis, oxidative phosphorylation, and fatty acid metabolism act as key regulators of macrophage specialization [13]. Murine studies suggest that M1 macrophages rely predominantly on glycolysis, whereas M2 macrophages depend on oxidative phosphorylation [14]. However, human macrophages appear to follow distinct metabolic programs, underscoring the risk of direct extrapolation from animal models and the need for investigation in clinically relevant systems [15, 16]. Building on this rationale, metabolic reprogramming has recently been explored as a means to modulate macrophage function [17]. The glycolytic regulator pyruvate kinase M2 (PKM2) has emerged as a central node in controlling macrophage polarization and effector functions. In murine models, pharmacological activation of PKM2 with compounds such as DASA-58 has been shown to shift macrophages toward an M2-like phenotype [18]. This immunometabolic regulation not only governs macrophage activation but also shapes their secretory activity [19].

Macrophage-derived extracellular vesicles (EVs), which carry proteins, lipids, and nucleic acids reflective of the polarization and metabolic state of their parental cells, have emerged as potent mediators of intercellular communication in tissue repair [20]. Through their molecular cargo, including microRNAs, they can modulate angiogenesis, osteogenesis, and extracellular matrix remodeling, extending the influence of macrophages beyond their immediate microenvironment [21]. However, most studies have focused on murine EVs or immortalized human macrophage lines, limiting their translational value [22]. A detailed characterization of EVs derived from primary human macrophages in the context of bone regeneration is therefore critically needed.

Hence, this study was designed to address these gaps by characterizing the metabolic states of primary human macrophage undergoing traditional M1 to M2 phenotypic shifts and then assessing the regenerative properties of the EVs derived from these unique metabolic and phenotypic states. Moreover, this study also explores whether DASA-58–mediated metabolic reprogramming of human macrophages can induce a phenotypic shift that promotes the release of regenerative EVs optimized for bone repair. Ultimately, this study provides the first functional validation of polarized human macrophage-derived EVs in angiogenesis and osteogenesis and demonstrates metabolic reprogramming can be harnessed to shift human macrophage phenotype and generate vesicles with hybrid properties tailored for tissue regeneration.

## 2. Results

### 2.1. Human macrophage polarization is associated with shifts in metabolic state

It is increasingly recognized that human macrophages exhibit unique metabolic characteristics to their mouse counterparts [23]. Therefore, we initially assessed the metabolic state of Mφ (naïve) human macrophages and investigated how this changes upon M1 (pro-inflammatory) and M2 (anti-inflammatory) macrophage polarization (Figure 1A). To validate successful human macrophage polarization into distinct phenotypes, we examined the mRNA expression of M1 markers such as *CXCL10* and *CD38*, both of which were significantly higher in M1 polarized macrophages compared to Mφ and M2 (p < 0.0001) (Figure 1B and C). Additionally, *VEGFA* mRNA expression was elevated in M1 compared to both Mφ (p < 0.0001) and M2 (p < 0.001) (Figure 1D). In contrast, the mRNA of *CD206* and *IL10*, characteristic M2 markers, were more highly expressed upon M2 polarization in comparison with their M1 counterparts (p < 0.01 and p < 0.001, respectively) (Figure 1E and F). Interestingly, while *IL10* mRNA expression was higher in M2 macrophages, ELISA results demonstrated increased IL-10 protein secretion in M1 macrophages compared to M2 (p < 0.001), suggesting post-transcriptional regulation in the M1 state (Figure 1G). Additionally, IL-8 and IL-6 protein expression levels were significantly elevated in M1 macrophages (p < 0.0001 and p < 0.05, respectively), supporting their pro-inflammatory phenotype (Figure 1H and I). Upon successfully validating human macrophage polarization, we used Agilent Seahorse technology to examine basal respiration rate and oxygen consumption rate (OCR) (Figure 1J). Basal respiration was significantly higher in M1 macrophages compared to Mφ (p < 0.01) and M2 macrophages (p < 0.0001), whereas M2 macrophages displayed a markedly lower basal respiration rate than Mφ (p < 0.001) (Figure 1K). A similar pattern was observed for overall OCR, with M1 macrophages exhibiting significantly higher values than both Mφ (p < 0.05) and M2 (p < 0.001) (Figure 1L). Next, to quantify glycolytic activity, we analyzed the mRNA expression of key glycolytic markers: *PKM2*, *HK2* (encoding hexokinase-2), and *HIF1A*. *PKM2* expression was significantly higher in M1 macrophages compared to both Mφ (p < 0.05) and M2 (p < 0.001), suggesting a glycolysis-driven metabolic profile (Figure 1M). Similarly, *HK2* expression was strongly upregulated in M1 macrophages (p < 0.0001) (Figure 1N), while *HIF1A* expression was also elevated in M1, reinforcing the shift toward aerobic glycolysis (Figure 1O). Collectively, these findings confirm that human M1 macrophages predominantly rely on glycolytic metabolism, exhibit high oxygen consumption, and secrete pro-inflammatory cytokines, while M2 macrophages maintain lower oxidative metabolism and glycolysis.

**Figure 1:**
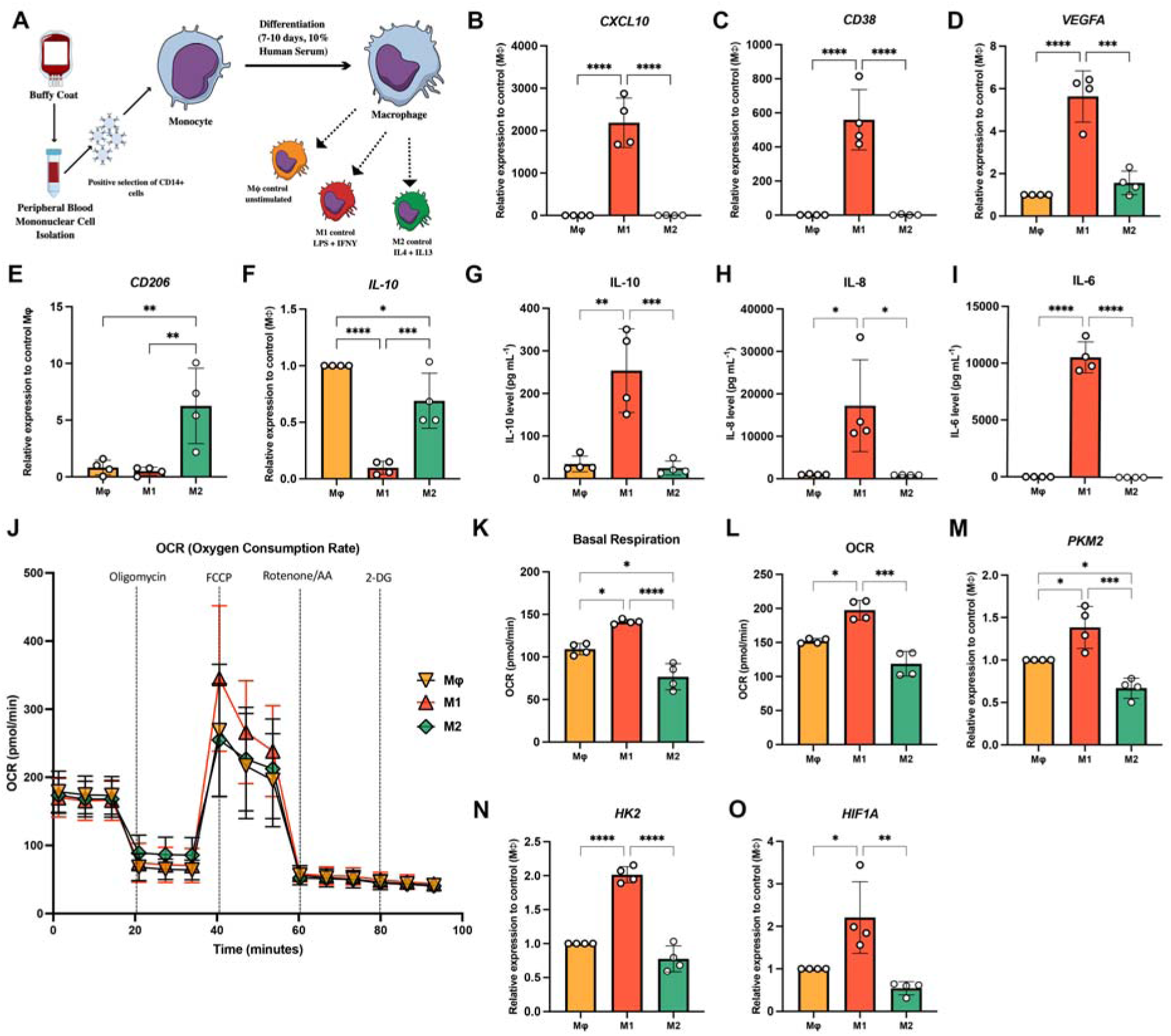
Human macrophage polarization is associated with changes in mRNA expression, cytokine levels and metabolic state. **(A)** Schematic illustration of human macrophage isolation and polarization. mRNA expression levels of M1 markers **(B)** *CXCL10* **(C)** *CD38* and **(D)** *VEGFA* and M2 markers **(E)** *CD206* and **(F)** *IL-10.* ELISA quantification of **(G)** IL-10, **(H)** IL-8 and **(I)** IL-6. **(J)** Seahorse plot of the Mitostress test of recording OCR in human macrophages following polarization to M1 and M2 at 3h. The values of the indicated respiratory parameters were calculated from the MitoStress test as **(K)** basal respiration and **(L)** oxygen consumption rate. mRNA expression levels of glycolytic markers **(M)** *PKM2*, **(N)** *HK2* and **(O)** *HIF1A.* Error bars in the graphs represent mean ± SD. N = 4 donors. ****p < 0.0001, ***p < 0.001, **p < 0.01, *p < 0.05 (one-way ANOVA and Tukey multiple comparison).

### 2.2. Human macrophages secrete EVs with a unique surface marker profile

As secretory cells, macrophages play an essential role in orchestrating adjacent cells during tissue repair through the release of paracrine factors often packaged within EVs [24]. Notably, EVs routinely reflect the characteristics of their parental cells; therefore, we examined whether EVs characteristics released from human macrophages differ according to polarization state, comparing EVs from unpolarized macrophages (EVs^Mφ^), M1-polarized macrophages (EVs^M1^), and M2-polarized macrophages (EVs^M2^). To investigate this, EVs were collected from the conditioned media (CM) of polarized macrophages by differential centrifugation. To ensure accurate characterization of EVs, we followed the MISEV guidelines [25]. Western blot analysis confirmed that GRP-94, an endoplasmic reticulum (ER) protein marker, was absent in all EV populations but was present in Mφ , M1, and M2 macrophages, confirming the purity of EV preparations (Figure 2A). In contrast, syntenin-1 and flotillin-1, two established EV protein markers, were consistently expressed in isolated EVs, verifying their identity. Transmission electron microscopy (TEM) further revealed EVs with the characteristic cup-shaped morphology, supporting their structural integrity (Fig.2B). To assess particle concentration, nanoparticle tracking analysis (NTA) was performed (Figure 2C), showing substantial donor-dependent variability which is expected for EVs derived from human primary cells [26]. This variability was particularly pronounced in EVs^M2^.

**Figure 2:**
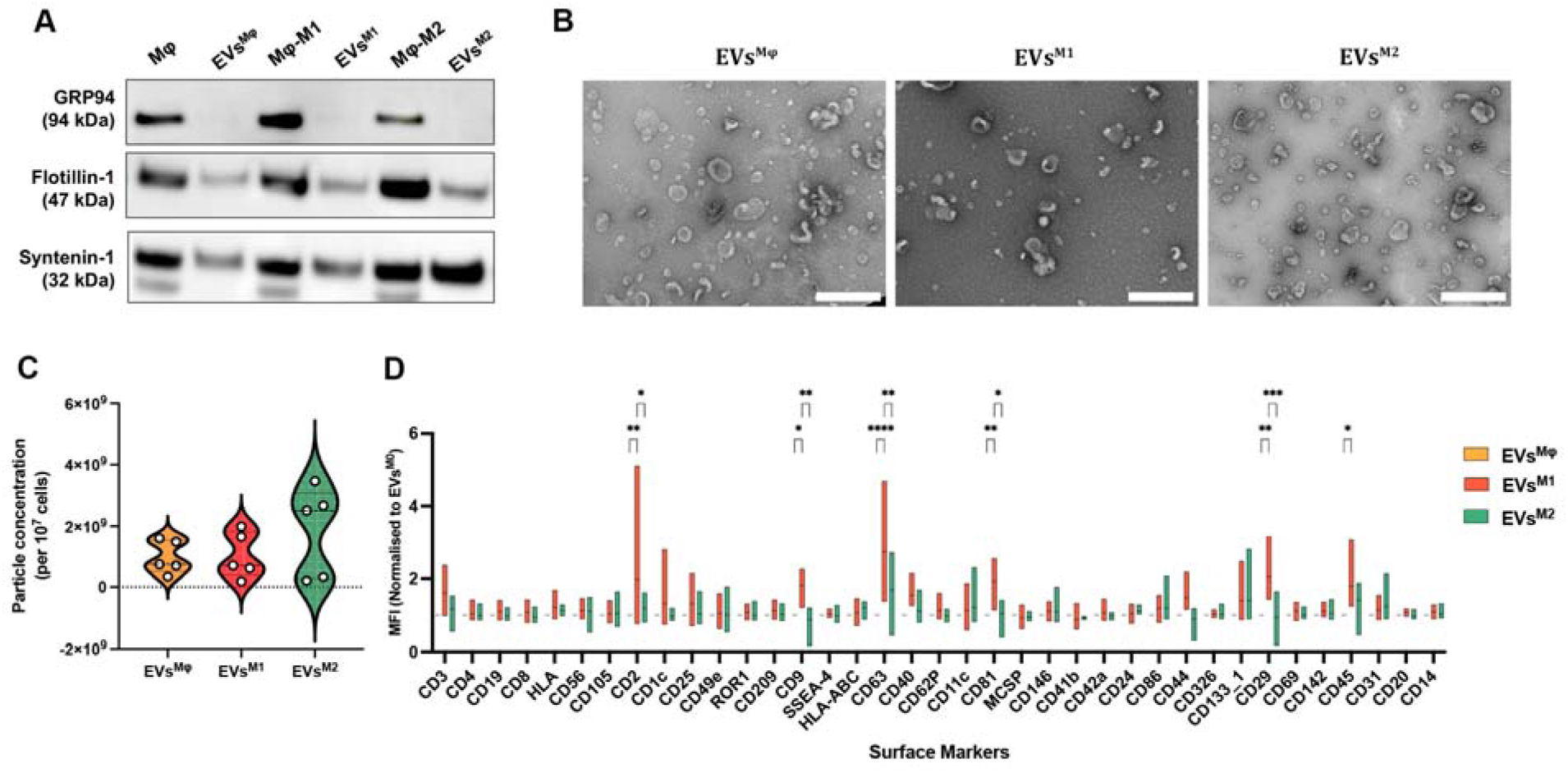
Human macrophages secrete EVs of similar morphology, quantity, but possess a unique surface marker profile depending on the polarization state. **(A)** Representative Western-blot analysis of non-EV marker GRP94 and, EV markers Flotilin-1 and Syntenin-1. **(B)** Representative transmission electron microscopy images of macrophage EVs. Scale bar=500 nm. **(C)** Quantitative analysis of EV concentration by Nanoparticle tracking analysis. N=5 donors. **(D)** Surface marker profiling of EVs with MACSplex Exosome Kit. ****p < 0.0001, ***p < 0.001, **p < 0.01, *p < 0.05 (two-way ANOVA and Tukey multiple comparison). N = 4 donors.

Given the distinct metabolic and functional differences between M1 and M2 macrophages, we next performed a detailed MACSPlex analysis to characterize the surface marker profiles of their EVs (Figure 2D). CD2 (cell adhesion molecule for immune cells) was significantly more abundant in EVs^M1^ compared to EVs^Mφ^ (p < 0.01) and EVs^M2^ (p < 0.05). A similar enrichment was observed for CD29 (ITGB1, cell surface receptor), which was predominantly expressed in EVs^M1^ relative to EVs^M2^ (p < 0.001). CD45 (lymphocyte common antigen) levels were also higher in EVs^M1^, but only when compared with EVs^Mφ^ (p < 0.05). The canonical tetraspanin markers, CD9, CD63, and CD81 were all significantly increased in EVs^M1^ compared with both EVs^Mφ^ and EVs^M2^, with no differences between EVs^Mφ^ and EVs^M2^. In contrast, EVs^M2^ showed a tendency toward higher expression of CD11c (monocyte marker), CD146 (melanoma cell adhesion molecule), CD133_1 (Prominin-1, membrane glycoprotein), and CD31 PECAM-1, cell adhesion molecule), indicating a relative enrichment of these markers in this subtype. Taken together, based on MISEV characterization guidelines, EVs released by human macrophages under different polarization states possess similar morphology and overall yield while consistently expressing canonical EV surface markers. However, their surface marker profiles undergo distinct changes depending on the polarization state.

### 2.3. Human macrophage-derived EVs ^M2^ but not EVs ^M1^ enhance angiogenesis

Angiogenesis is a critical component of bone fracture repair, as endothelial cells play a key role in forming new blood vessels to support tissue regeneration [27]. To investigate the role of human macrophage-derived EVs in this process, we evaluated their impact on different aspects of angiogenesis, including endothelial cell proliferation, migration and tube formation.

Firstly, we assessed human umbilical vein endothelial cell (HUVEC) proliferation rates following 1 mg/mL EV treatment. While both EVs^M1^ and EVs^M2^ significantly increased HUVEC proliferation compared to endothelial cell basal media (EBM) controls (p < 0.01 and p < 0.001, respectively), no difference was observed with EVs^Mφ^ and EBM controls (Figure 3A). To further explore the contribution of macrophage-derived EVs to wound healing and angiogenesis, we performed a HUVEC scratch assay to assess migration (Figure 3B, C and D). Following the scratch, 1 µg/mL of EVs were added, and scratch closure was monitored over a 24-hour period (Figure 3D). To capture the most distinct differences between groups, migration was quantified at a 12-hour time point (Figure 3B). Interestingly, only EVs^M2^ significantly enhanced HUVEC migration compared to all other groups, while EVs^Mφ^ and EVs^M1^ failed to induce migration, showing no significant difference from the EBM control. Moreover, EVs^M1^-treated cells exhibited significantly lower migration rates compared to EVs^M2^ (p < 0.0001), further confirming the unique potency of EVs^M2^.

**Figure 3:**
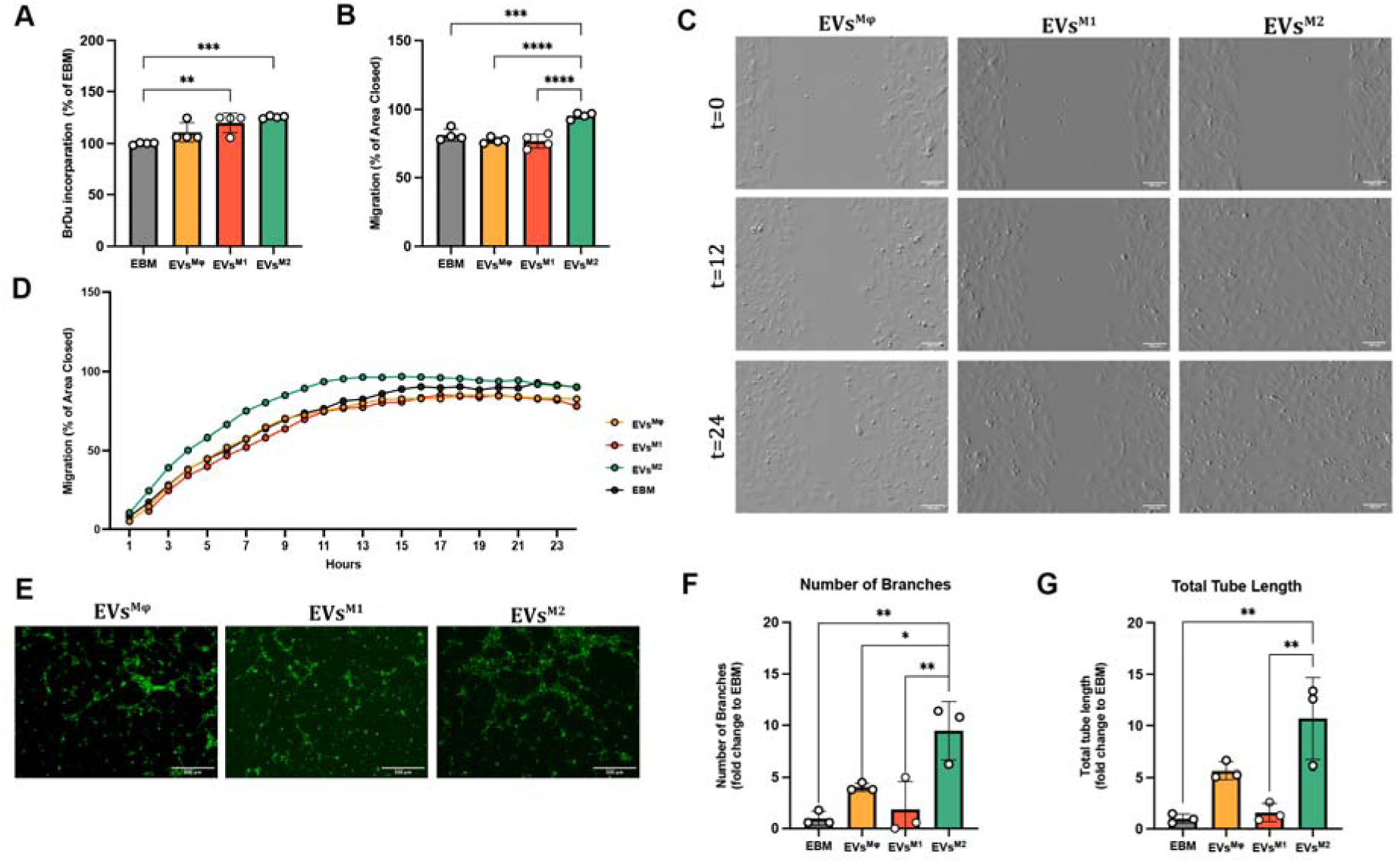
Human macrophage-derived EVs differentially regulate endothelial angiogenesis with EVs^M2^ promoting migration and tube formation. **(A)** Quantitative analysis of BrdU incorporation assay measuring endothelial cell proliferation. **(B)** Quantitative analysis of HUVEC migration and **(C)** representative images of HUVEC migration. Scale bar=100 µm. **(D)** Plot of HUVEC migration for 24 hours. **(E)** Representative images of GFP-HUVECs which were treated with 1 µg/mL EVs for 9 hours on GelTrex. Scale bar=500 µm. **(F)** Quantitative analysis of number of branches and **(G)** total tube length. Error bars in the graphs represent mean ± SD of EVs obtained from three or four independent donors. ****p < 0.0001, ***p <0.001, **p < 0.01, *p <0.05 (one-way ANOVA and Tukey multiple comparison).

To determine whether these effects on HUVEC proliferation and migration were potentially linked to the tube formation capacity of endothelial cells, 1 µg/mL of EVs were added to GFP-HUVECs and angiogenesis was tracked and the number of branches and total tube length quantified. Both EVs^Mφ^ and EVs^M2^ significantly enhanced the formation of tube-like structures (Figure 3E) when compared to untreated control, demonstrating their pro-angiogenic potential. Quantitative analysis revealed that EVs^M2^ treatment led to the most robust and significant increase in branch formation, with EVs^M2^-treated endothelial cells significantly outperforming EVs^Mφ^ (p < 0.01) and EVs^M1^ (p < 0.05) (Figure 3F). In contrast, EVs^M1^ had no significant effect on tube formation compared to the negative control (EBM), indicating a lack of pro-angiogenic properties. This was further verified when quantifying total tube length, where EVs^M2^ treatment significantly outperformed EVs^M1^ (p < 0.01) (Figure 3G), demonstrating the angiogenic properties of macrophage derived EVs is strongly dependent on the parent cell phenotype.

Across all three angiogenesis assays, EVs^M2^ demonstrated superior activity, enhancing tube formation, proliferation, and migration, thereby reinforcing their pro-angiogenic potential. In contrast, EVs^M1^ showed no significant effect on HUVEC tube formation or migration, suggesting that human macrophage-derived EVs^M1^ have a limited role in angiogenesis.

### 2.4. Human macrophage-derived EVs^M1^ but not EVs ^M2^ enhance human MSC migration and early osteogenesis

Given the crucial role of the macrophage secretome in mediating early-stage bone healing [4], we next investigated the effect of human macrophage-derived EVs on human mesenchymal stromal cell (MSC) proliferation, migration, and osteogenic differentiation. Firstly, to examine the effect of EVs on MSC proliferation, a BrdU incorporation assay was performed using 1 µg/mL of EVs. All EV groups significantly increased proliferation compared to EBM (p < 0.001), with EVs^M1^ showing the most pronounced effect (Figure 4A). Next, to assess MSC migration a scratch assay was performed. After scratching, 1 µg/mL of EVs were added, and closure was monitored over 24 hours (Figures 4B, C and D). As full scratch closure was observed in all groups by 24 hours (Figure 4C and D), we analysed an earlier time point (12 hours), consistent with the HUVEC assay conditions. At this time point, EVs^M1^ significantly increased MSC migration compared to both the negative control and EVs^Mφ^, indicating that EVs^M1^ promoted a more rapid MSC migration (p < 0.01) (Figure 4B).

**Figure 4:**
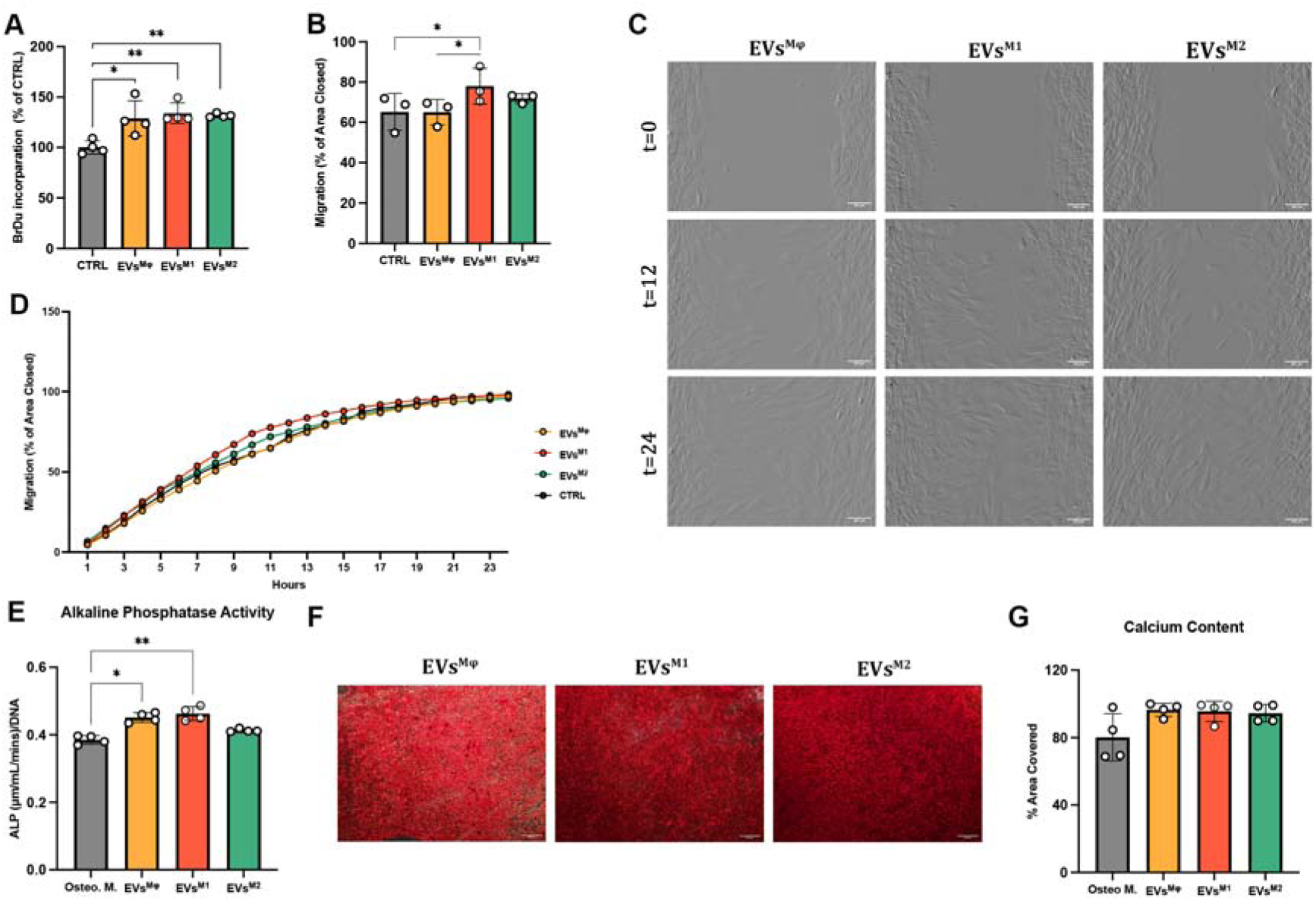
EVs^M1^ enhance MSC proliferation, migration and early osteogenesis but demonstrate no differences in late-stage mineralization. **(A)** Quantitative analysis of BrdU incorporation assay measuring MSC proliferation. **(B)** Quantitative analysis of MSC migration and **(C)** representative images of MSC migration. Scale bar=100 µm. **(D)** Plot of MSC migration for 24 hours and **(E)** alkaline phosphatase activity at day 7. **(F)** Alizarin Red staining at day 14 for mineralized matrix deposition (Scale bar=100 µm) and **(G)** quantification of calcium content. Error bars in the graphs represent mean ± SD of EVs obtained from three or four independent donors. ****p < 0.0001, ***p < 0.001, **p < 0.01, *p < 0.05 (one-way ANOVA and Tukey multiple comparison).

To investigate the effects on osteogenesis, EVs were added to MSCs every three days at a concentration of 1 µg/mL, and Alkaline Phosphatase Activity (ALP) was measured on day 7. The results showed that EVs^Mφ^ and EVs^M1^ significantly increased ALP activity compared to control osteogenic media (p < 0.01), with EVs^M1^ once again demonstrating the most pronounced effect, whilst no significant differences in ALP activity were observed with EVs^M2^ (Figure 4E). To further examine osteogenic differentiation, Alizarin Red staining was performed on day 14 to assess mineral deposition (Figure 4F). Quantification of calcium content revealed no significant differences between groups, suggesting that while EVs^M1^ enhanced early-stage osteogenesis, no differences were observed at later stages among MSCs treated with human macrophage-derived EVs (Figure 4G).

### 2.5. Shifts in macrophage phenotype, particularly M1 polarization, modulates the miRNA cargo of EVs

Given the phenotype-specific differences in the biological effects of macrophage-derived EVs on angiogenesis and osteogenesis, we next investigated whether this may be mediated by shifts in their miRNA cargo. Absolute concentrations of miRNAs detected in the Next Generation Sequencing (NGS) data were calculated using miND® spike-ins [28]. First, to identify the most abundant miRNAs, we calculated the relative distribution of miRNAs in EVs^Mφ^, EVs^M1^, and EVs^M2^. Although minor changes in proportions were observed, the same 15 miRNAs were most abundant across all groups (Figure 5A). Consequently, we focused on differentially expressed (DE) miRNAs. In EVs^M1^ vs. EVs^Mφ^, 12 miRNAs were upregulated and 2 downregulated. Among the upregulated set were inflammation-related miRNAs such as hsa-miR-155-5p, hsa-miR-146b-5p, and hsa-miR-146a-5p, indicating that upon M1 polarization a pro-inflammatory signature was also present in the EV cargo at a miRNA level [29]. In contrast, hsa-miR-484 and hsa-miR-1307-3p were downregulated in EVs^M1^, compared to EVs^Mφ^ (Figure 5B). Notably, only one miRNA, hsa-miR-484, was downregulated in EVs^M2^, with higher expression found in EVs^Mφ^ (Figure 5C). When comparing EVs^M1^ to EVs^M2^, inflammation-related miRNAs again appeared among the most upregulated, while hsa-miR-1307-3p was downregulated, suggesting that lower levels of this miRNA is associated with M1 polarization. Conversely, hsa-miR-423-3p and hsa-miR-133a-3p were more abundant in EVs^M2^ (Figure 5D). Normalizing DEs miRNA expression to their Mφ counterparts, principal component analysis (PCA) revealed two distinct profiles for EVs^M1^ and EVs^M2^, indicating that polarization regulates the miRNA content of EVs (Figure 5E). To identify miRNAs associated with polarization, we analyzed DE miRNA patterns across all groups (Figure 5F). While most differences were detected between EVs^M1^ and EVs^Mφ^, some miRNAs were specifically regulated by M1 or M2 polarization, whereas hsa-miR-484 was characteristic of EVs^Mφ^. Heatmap visualization further highlighted donor-to-donor variability in miRNA expression patterns (Figure 5G).

**Figure 5:**
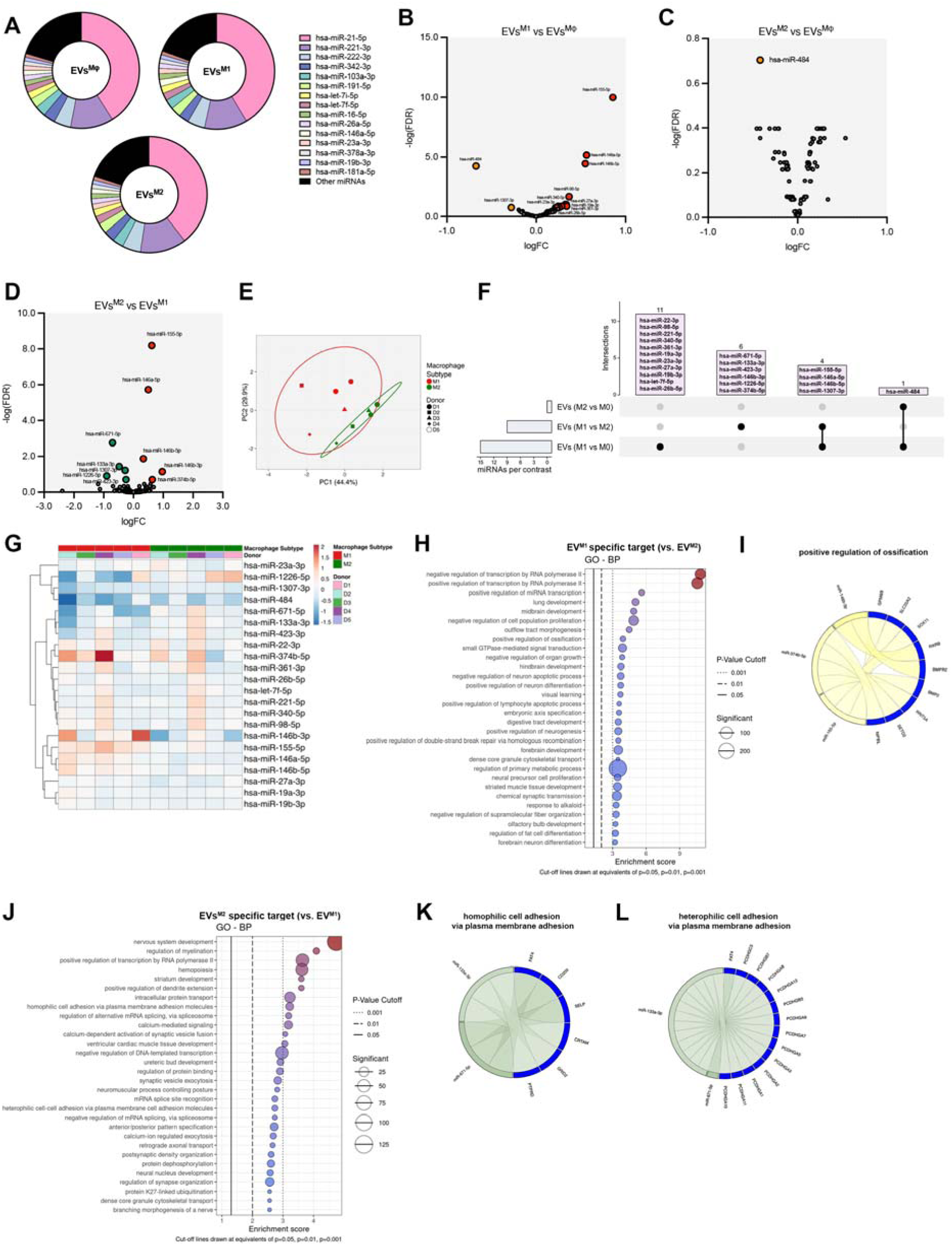
EVs^M1^ display a distinct miRNA profile compared to EVs^Mφ^ and EVs^M2^. **(A)** Relative distribution of miRNAs in EVs^Mφ^, EVs^M1^, and EVs^M2^. Pie charts depict the total miRNA pool and highlight the 15 most abundant miRNAs in each group. Volcano plots illustrate differentially expressed (DE) miRNAs between **(B)** EVs^M1^ vs. EVs^Mφ^, **(C)** EVs^M2^ vs. EVs^Mφ^, and **(D)** EVs^M1^ vs. EVs^M2^. Statistical significance was defined as a false discovery rate (FDR) < 0.2. **(E)** Principal component analysis (PCA) of DE miRNAs, with each donor represented by a distinct symbol. **(F)** Upset plot summarizing the overlap of DE miRNAs across comparisons. **(G)** Heatmap of DE miRNAs, showing expression patterns between groups. **(H)** Top 30 significantly enriched *GO* biological processes associated with upregulated miRNA target genes in EVs^M1^ vs EVs^M2^ (EVs^M1^ specific miRNA subset). **(I)** Chord diagram showing the involvement of miRNAs in the positive regulation of ossification. **(J)** Top 30 significantly enriched GO biological processes associated downregulated miRNA target genes in EVs^M1^ vs EVs^M2^ (EVs^M2^ specific miRNA subset). **(I)** Chord diagram showing the involvement of miRNAs in **(K)** homophilic cell adhesion via plasma membrane adhesion and **(L)** heterophilic cell adhesion via plasma membrane adhesion. N=5 donors.

Functional gene ontology (GO) analysis of miRNA targets genes from EV associated miRNAs identified several biological processes (BP) involved in transcriptional regulation, developmental process and regenerative functions. When comparing differently upregulated (higher abundant in EVs^M1^) and downregulated (higher abundant in EVs^M2^) miRNAs between EVs^M1^ versus EVs^M2^, different enriched BPs were found based on the miRNA target gene subsets for EVs^M1^ and EVs^M2^. Among the top 30 leading biological processes associated with higher abundant miRNAs in EVs^M1^ were linked to “regulation of transcription by RNA polymerase II” (GO:0000122, GO:0045944) and development processes (GO:0030324, GO:0030901) (Figure 5H). Of note, GO enrichment analysis identified “positive regulation of ossification” (GO:0045778) (Figure 5I), among the top 30 enriched biological processes, which is consistent with our findings of increased ALP activity in Figure 4. In comparison, the top 30 biological process terms for EVs^M2^ were linked to intercellular protein transport (GO:0006886), regulation of protein binding (GO:0043393), and cell adhesion molecules, specifically homophilic and heterophilic cell adhesion mediated by the plasma membrane (GO:0007156, GO:0007157) (Figure 5J,5K and 5L). These processes are fundamental for angiogenesis, supporting our findings that EVs^M2^ promote pro-angiogenic signaling. Additional results for GO molecular function (MF) terms showed that miRNAs targets associated with EVs^M1^ were linked to nuclear receptor binding (GO:0016922) and activity (GO:0004879) functions involved in osteogenic regulation, whereas EVs^M2^ showed enrichment in cadherin binding (GO:0045296), consistent with roles in cell adhesion (Supplementary Figure 5A and B). Taken together, these findings suggest that EVs^M1^ are primarily associated with osteogenesis, whereas EVs^M2^ are more closely linked to mechanisms supporting angiogenesis.

### 2.6. DASA-58 modulates human macrophage metabolism, resulting in a balanced hybrid phenotype with both pro-inflammatory and anti-inflammatory features

Given the distinct roles of human EVs^M1^ and EVs^M2^ in modulating angiogenesis and osteogenesis respectively, we next sought to generate a hybrid macrophage phenotype to harness the benefits of both polarization states. To achieve this and given that we have previously demonstrated that human macrophage phenotype is associated with distinct metabolic states, we examined the effect of the PKM2 glycolytic enzyme activator DASA-58, as it has been shown to induce metabolic reprogramming in murine macrophages [18]. However, given the known metabolic differences between murine and human macrophages [12], we first evaluated two concentrations of DASA-58 (50 µM and 100 µM) in human macrophages. A preliminary cytotoxicity assay confirmed that both concentrations of DASA-58 were well tolerated by the human macrophages, with no detectable cytotoxic effects (Supplementary Figure 2A and B). Subsequently, macrophages were pretreated with DASA-58 for 2 hours before polarization to M1 phenotype to induce an inflammatory reaction consistent with the initial stages of healing (Figure 6A).

**Figure 6:**
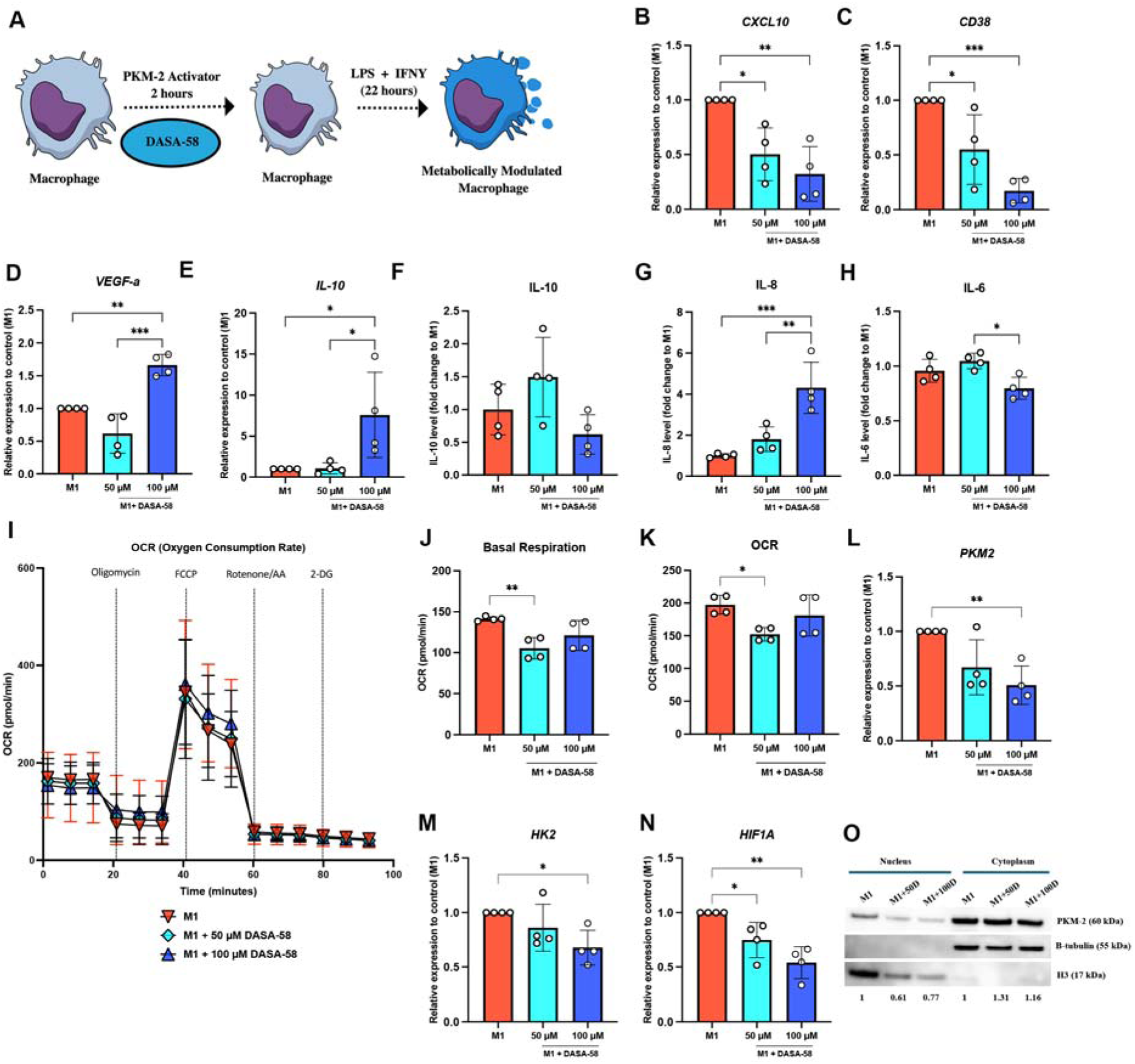
DASA-58 modulates macrophage metabolism and polarization, shifting M1 macrophages toward an M2-like phenotype. **(A)** Schematic illustration of PKM2 activator DASA-58 treatment on human macrophages. mRNA expression levels of M1 markers **(B)** *CXCL10*, **(C)** *CD38* and **(D)** *VEGFA* and M2 marker **(E)** *IL-10.* ELISA quantification of **(F)** IL-10, **(G)** IL-8 and **(H)** IL-6. **(I)** Seahorse plot of the Mitostress test of recording OCR in human macrophages following DASA-58 treatment at 3h. The values of the indicated respiratory parameters were calculated from the MitoStress test **(J)** basal respiration and **(K)** as oxygen consumption rate. mRNA expression levels of glycolytic markers **(L)** *PKM2*, **(M)** *HK2* and **(N)** *HIF1A.* **(O)** Western blot analysis of PKM2, Histone H3, and Tubulin in cytoplasmic and nuclear fractions in human macrophages following DASA-58 treatment. Histone H3 was used as a nuclear marker, and Tubulin was used as a cytoplasmic marker to confirm the purity of subcellular fractions. Error bars in the graphs represent mean ± SD. N = 4 donors. ****p < 0.0001, ***p <0.001, **p < 0.01, *p <0.05 (one-way ANOVA and Tukey multiple comparison).

To investigate the phenotypic impact of DASA-58 mediated metabolic reprogramming of M1 macrophages, we analyzed the expression of M1 polarization markers *CXCL10*, *CD38*, and *VEGFA*. Interestingly, DASA-58 exhibited a dose-dependent effect, with 100 µM DASA-58 significantly reducing *CXCL10* (p < 0.01) and *CD38* (p < 0.01) expression compared to M1 macrophages (Figure 6B and C). Surprisingly, DASA-58 also increased *VEGFA* expression, indicating an enhanced angiogenic response following treatment (p < 0.01) (Figure 6D). Another notable finding was the upregulation of *IL10*, an important marker of human M2 macrophages, which showed significantly higher mRNA expression levels exclusively at 100 µM DASA-58 (p < 0.05) (Figure 6E). To further assess cytokine secretion, we measured IL-10, IL-6, and IL-8 protein levels using ELISA. No significant changes were observed in IL-10 and IL-6 protein levels following treatment with 100 µM DASA-58 (Figure 6F and G). However, IL-8 secretion was significantly increased at 100 µM (p < 0.001), demonstrating a dose-dependent response (Figure 6H).

To verify that DASA-58 can modulate metabolism in human macrophages, we measured basal respiration and OCR in polarized macrophages using Seahorse analysis (Figure 6I). As demonstrated previously (Figure 1K), basal respiration was significantly higher in M1 macrophages compared to M2. Interestingly, treatment with 50 µM DASA-58, and to a lesser extent with 100 µM, significantly reduced basal respiration compared to M1 macrophages (p < 0.05), reaching a level closer to that seen in M2 macrophages (Figure 6J). A similar trend was observed in OCR measurements, indicating that DASA-58 altered mitochondrial activity in a dose-dependent manner (Figure 6K). To determine whether DASA-58 affected glycolytic mRNA expression, we measured mRNA levels of *PKM2*, *HK2*, and *HIF1A*. Treatment with 100 µM DASA-58 significantly decreased *PKM2* (p < 0.01), *HK2* (p < 0.05), and *HIF1A* (p < 0.01) expression compared to M1 macrophages, indicating a suppression of glycolytic mRNA activity in macrophages exposed to the DASA-58 (Figure 6L, M and N).

Lastly, to examine whether DASA-58 influenced PKM2 colocalization, we analyzed its nuclear and cytoplasmic expression [30]. The results showed that DASA-58 treatment reduced nuclear PKM2 levels compared to M1 macrophages, suggesting a shift in PKM2, as described before (Figure 6O).

Overall, these results indicate that PKM-2 activation in human macrophages alters the M1 phenotype by modulating metabolism, mRNA expression, and cytokine secretion toward a more M2-like hybrid state. This suggests that targeting PKM-2 with DASA-58 can induce a unique macrophage phenotype with hybrid features of M1 and M2 states in a dose-dependent manner.

### 2.7. DASA-58 reprograms M1 macrophages to produce EVs with enhanced angiogenic activity without compromising osteogenic potential

Given that DASA-58 treatment induces a metabolically modulated, hybrid human macrophage phenotype, and that EVs inherit key characteristics of their parental cells [31], we next investigated the properties of EVs derived from these reprogrammed macrophages. Before assessing their functional effects, we first characterized the EVs from metabolically modulated macrophages.

Following standard characterization based on MISEV guidelines using WB, NTA and TEM, no significant differences were observed between EVs derived from macrophages treated with 50 µM and 100 µM DASA-58 (to be known as EVs^50D^ and EVs^100D^ respectively) (Supplementary Figure 3A, B and C). However, MACSPlex analysis revealed a significantly higher expression of fibronectin receptor CD49E in EVs^100D^ compared to EVs^M1^ and EVs^50D^ (p < 0.01 and p < 0.05, respectively) (Supplementary Figure 3D). To determine whether the metabolic modulation induced by DASA-58 affects the biological function of EVs, we examined the effects of EVs^50D^ and EVs^100D^ on angiogenesis and osteogenesis, using EVs^M1^ as a control.

Firstly, we examined the effects of these EVs on HUVEC proliferation. Both EVs^50D^ and EVs^100D^ significantly promoted HUVEC proliferation compared to EBM control (p < 0.0001) (Figure 7A). To assess their impact on migration, we performed a scratch assay (Figure 7B and C). While EVs^50D^ induced a response similar to EVs^M1^ (Supplementary Figure 4A), only EVs^100D^ significantly increased HUVEC migration compared to both EBM and EVs^M1^ (p < 0.05 and p < 0.01, respectively) (Figure 7B). In a tube formation assay, EVs^100D^ significantly increased the number of branches (p < 0.01), and total tube length (p < 0.001) compared to EVs^M1^ (Figure 7D, E and F). EVs^50D^ also enhanced tube formation, but with a lower effect size, increasing the number of branches (p < 0.05) while showing a similar trend to EVs^100D^ for total tube length (Figure 7E and F). Both EVs^50D^ and EVs^100D^ therefore enhanced angiogenesis, demonstrating functional differences from EVs^M1^ due to metabolic modulation. Notably, EVs^100D^ promoted angiogenesis to levels comparable with EVs^M2^, effectively overcoming the angiogenic limitations of EVs^M1^. Together, these findings show that PKM2-driven metabolic modulation enables human macrophages to generate EVs with pro-angiogenic properties comparable to EVs from M2 macrophages, which are preserved even after M1 polarization.

**Figure 7:**
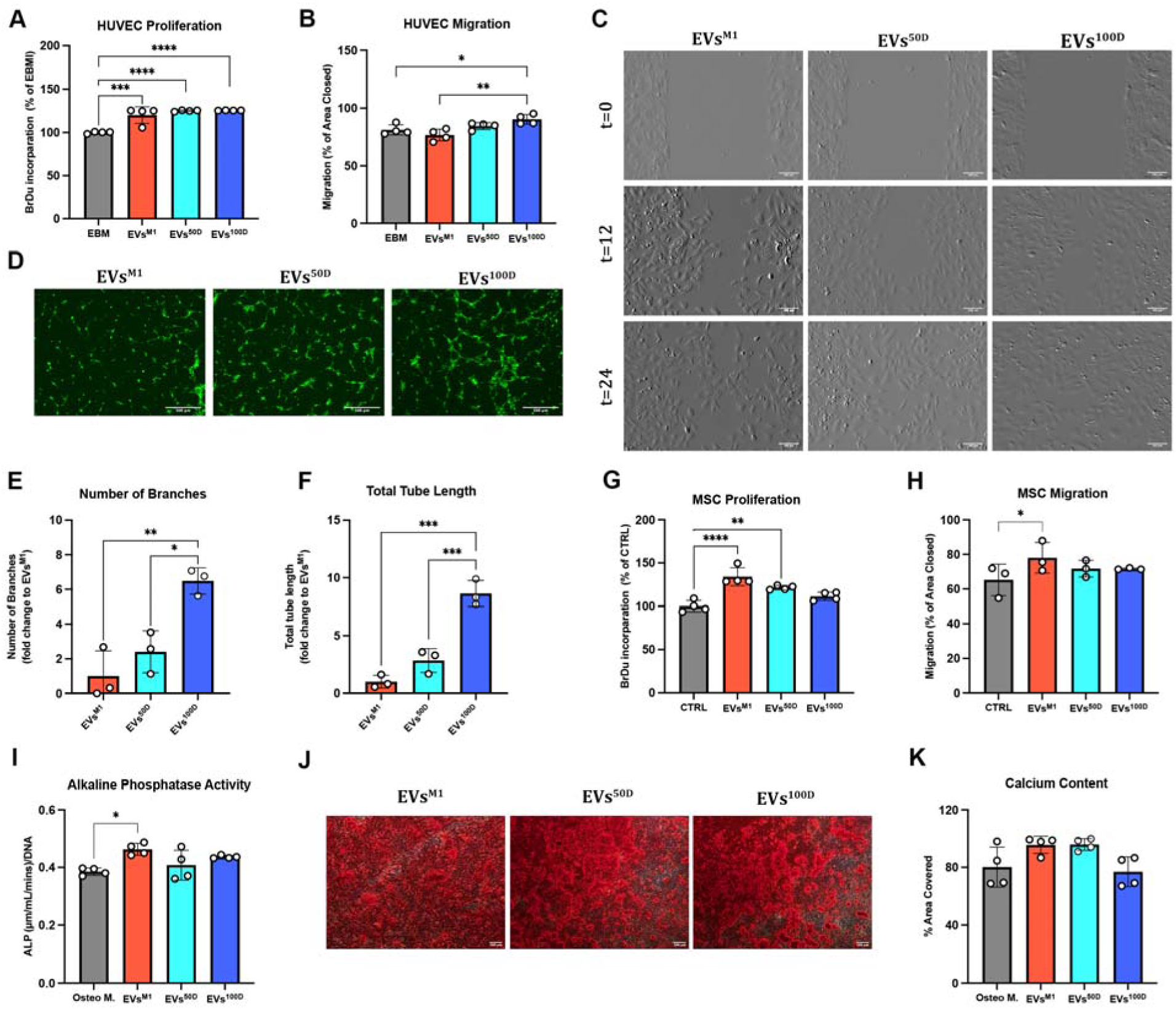
EVs Derived from Metabolically Modulated M1 Macrophages Exhibit Distinct Angiogenic and Osteogenic Effects. **(A)** Quantitative analysis of BrdU incorporation assay measuring endothelial cell proliferation. **(B)** Quantitative analysis of HUVEC migration and **(C)** representative images of HUVEC migration. Scale bar= 100 µm. **(D)** Representative images of GFP-HUVECs which were treated with 1 µg/mL EVs for 9 hours on GelTrex. Scale bar=500 µm. (**E**) Quantitative analysis of number of branches and (**F**) total tube length. **(G)** Quantitative analysis of BrdU incorporation assay measuring MSC proliferation. **(H)** Quantitative analysis of MSC migration. **(I)** Alkaline phosphatase activity at day 7. **(J)** Alizarin Red staining at day 14 for mineralized matrix deposition (Scale bar=100 µm) and **(K)** quantification of calcium content. Error bars in the graphs represent mean ± SD of EVs obtained from three or four independent donors. ****p < 0.0001, ***p < 0.001, **p < 0.01, *p < 0.05 (one-way ANOVA and Tukey multiple comparison). Error bars in the graphs represent mean ± SD of EVs obtained from three or four independent donors.

Given that we have shown that EVs^M1^ were more effective in supporting osteogenesis, we next sought to determine whether metabolically modulated macrophages polarized under M1 conditions after the DASA-58 treatment maintained these pro-osteogenic properties. Initially assessing the impact of these EVs on MSC proliferation (Figure 7G), our findings revealed that EVs^50D^ significantly increased MSC proliferation compared to the M1 control (p < 0.01). To further evaluate their role in osteogenesis, we analyzed MSC migration (Supplementary Figure 4B and C). Interestingly, both EVs^50D^ and EVs^100D^ exhibited a similar trend to the negative control (Figure 7H), showing no significant effect on MSC migration but also no reduction compared to EVs^M1^, indicating an unchanged migratory potential. This trend was further confirmed by early ALP activity at day 7, where EVs^50D^ significantly decreased ALP activity (p < 0.05) compared to EVs^M1^, while EVs^100D^ exhibited no change, maintaining a level similar to EVs^M1^ (Figure 7I). Further validation using Alizarin Red staining revealed comparable levels of mineralization among all groups (Figure 7J). However, calcium deposition at day 14 was significantly reduced in EVs^100D^ treated cells compared to EVs^M1^ and EVs^50D^ (p < 0.05), but no differences were observed when compared to the standard osteogenic differentiation medium (Figure 7K). Taken together, these findings indicate that metabolic modulation of M1-primed human macrophages results in the release of EVs with an unchanged impact on osteogenesis.

### 2.8. EVs^100D^ Exhibit a Hybrid miRNA Cargo Between M1 and M2 Phenotypes

To investigate whether changes in EV regenerative properties seen following metabolic modulation may be mediated by changes in EV cargo, we next investigated whether DASA-58 modulation of macrophages altered the EV miRNA profile. Since most changes were observed in the biological activity of EVs^100D^ in comparison with EVs^50D^, we focused on characterizing the EV^100D^ cargo and determining whether this profile resembled an intermediate state between EVs^M1^ and EVs^M2^. To first characterize the EVs^100D^ cargo, we examined the most abundant miRNAs. The top 14 miRNAs were largely shared with EVs^Mφ^, EVs^M1^, and EVs^M2^, with the exception that EVs^100D^ lacked hsa-miR-181a-5p in the top 15 miRNA list and instead was enriched in hsa-miR-423-3p, a miRNA we previously shown to be upregulated in EVs^M2^ compared with EVs^M1^ (Figure 8A). Differential expression analysis between EVs^100D^ and EVs^M1^ identified 14 upregulated and 16 downregulated miRNAs. Among the downregulated set were inflammation-related miRNAs, including hsa-miR-146b-5p and hsa-miR-146a-5p, suggesting that EVs^100D^ exhibited a reduced pro-inflammatory cargo relative to EVs^M1^, while still showing strong regulation of hsa-miR-423-3p (Figure 8B). Conversely, when EVs^100D^ were compared with EVs^M2^, inflammation-related miRNAs such as hsa-miR-155-5p and hsa-miR-374b-5p were enriched (Figure 8C). These results suggest that the miRNA cargo of EVs^100D^ contains elements of both EVs^M1^ and EVs^M2^.

**Figure 8:**
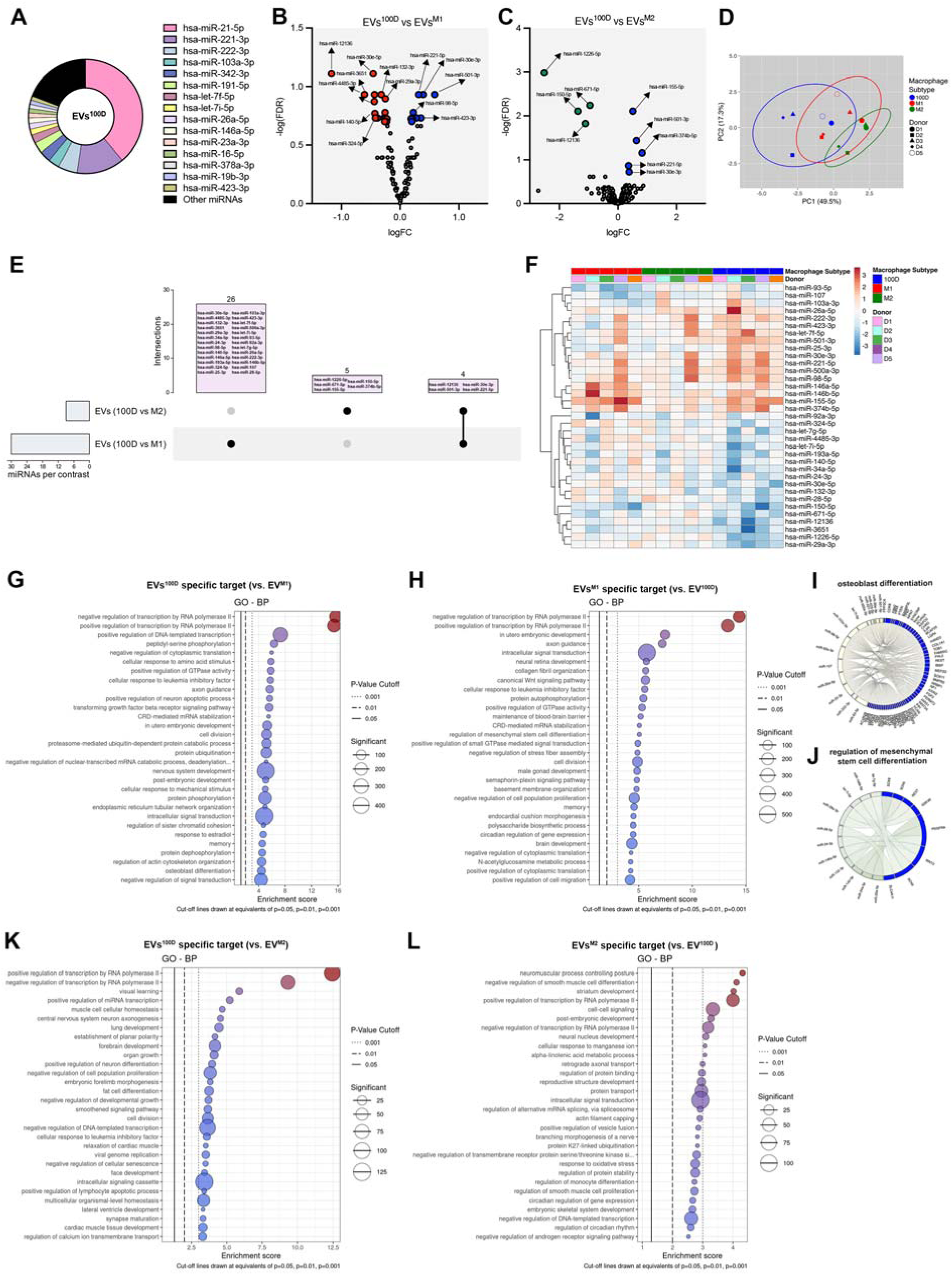
**(A)** Relative distribution of miRNAs in EVs^100D^, pie chart depicts the total miRNA pool and highlight the 15 most abundant miRNAs in each group. Volcano plots illustrate differentially expressed (DE) miRNAs between **(B)** EVs^100D^ vs. EVs^M1^ and **(C)** EVs^100D^ vs. EVs^M2^. Statistical significance was defined as a false discovery rate (FDR) < 0.2. **(D)** Principal component analysis (PCA) of DE miRNAs, with each donor represented by a distinct symbol. **(E)** Upset plot summarizing the overlap of DE miRNAs across comparisons. **(F)** Heatmap of DE miRNAs, showing expression patterns between groups. N=5 donors. Top 30 significantly enriched GO biological processes associated with upregulated miRNA target genes in **(G)** EVs^100D^ vs EVs^M1^ (EVs^100D^ specific miRNA subset) and **(H)** EVs^M1^ vs EVs^100D^ (EVs^M1^ specific miRNA subset). Chord diagram showing the involvement of upregulated miRNAs in EVs^100D^ **(I)** in osteoblast differentiation, and in EVs^M1^ **(J)** in the positive regulation of mesenchymal stem cell differentiation. Top 30 significantly enriched GO biological processes associated with upregulated miRNA target genes in **(K)** EVs^100D^ vs EVs^M2^ (EVs^100D^ specific miRNA subset) and **(L)** EVs^M2^ vs EVs^100D^ (EVs^M2^ specific miRNA subset). N=5 donors.

PCA demonstrated that EVs^100D^ overlapped with both EVs^M1^ and EVs^M2^ in terms of differentially expressed miRNAs, indicating that DASA-58 drives the generation of an intermediate EV profile (Figure 8D). Interestingly, an Upset plot revealed considerable differences in cargo between EVs^100D^ and EVs^M1^, despite both EV populations being derived from macrophages stimulated with LPS and IFN-γ. Notably, four miRNAs were differentially expressed across EVs^M1^ and EVs^M2^ in relation to EVs^100D^: hsa-miR-501-3p, hsa-miR-30e-3p, and hsa-miR-221-5p were upregulated, while hsa-miR-12136 was downregulated, highlighting that DASA-58 treatment induced a unique EV cargo signature in EVs^100D^ (Figure 8E). Heatmap analysis further supported this, showing that some miRNAs in EVs^100D^ mirrored trends observed in EVs^M1^, while others aligned with EVs^M2^ patterns, with additional variability attributed to the EVs donor (Figure 8F). GO enrichment analysis comparing miRNA target genes from higher abundant miRNAs in EVs^100D^ in compared to EVs^M1^ revealed associations primarily related to the regulation of transcription (GO:0000122, GO:0045944, GO:0045893), whereas the target genes of miRNAs with a higher expression in EVs^M1^ compared to EVs^100D^ showed overlapping top associations in “transcription by RNA polymerase II” (GO:0000122, GO:0045944) (Figure 8G and H). Interestingly, in these top 30 miRNA targets, EVs^100D^ showed a connection to biological processes related to “osteoblast differentiation” (GO:0001649), whereas EVs^M1^ were linked to “regulation of mesenchymal stem cell differentiation” (GO:2000739), potentially explaining their role in initiating endochondral ossification (Figure 8I and J). This regulatory shift toward osteoblast differentiation in EVs^100D^ may occur as inflammation decreases. In contrast, comparisons between EVs^100D^ and EVs^M2^ did not yield significant biological processes directly related to angiogenesis or osteogenesis (Figure 8K and L). Interestingly, further GO-MF results revealed that upregulated miRNAs in EVs^100D^ in comparison to EVs^M2^ were linked to cadherin binding (GO:0045296), similar to EVs^M2^ when it was compared with EVs^M1^, supporting a role for cell adhesion in angiogenesis (Supplementary Figure 5C and 5D). Moreover, when EVs^100D^ vs EVs^M2^ were compared, EVs^100D^ displayed nuclear receptor binding (GO:0016922) and activity (GO:0004879), paralleling profiles observed in EVs^M1^ and consistent with osteogenic regulation (Supplementary Figure 5E and 5F). Collectively, the findings suggest that EVs^100D^ possess a distinct miRNA cargo that integrates features from both EVs^M1^ and EVs^M2^, thereby enhancing their therapeutic potential for regeneration.

## 3. Discussion

Macrophages play a central role in repair, orchestrating the transition from inflammation to regeneration through dynamic phenotypic changes. Although traditionally described within a binary M1/M2 framework, this classification is overly simplistic and fails to capture the intermediate and overlapping states needed to contribute to tissue repair [32]. Macrophage polarization is closely coupled to cellular metabolism, with metabolic rewiring shaping not only their functional specialization but also their secretory activity. While murine studies have established associations between glycolysis, oxidative phosphorylation, and macrophage phenotype, the metabolic programs of human macrophages remain poorly defined [14, 33]. These differences are particularly relevant for their secretomes, especially EVs, which have emerged as potent mediators of macrophage function. EVs mirror many of the functional features of their parental cells and have been proposed both as therapeutic targets and as potential therapeutic agents [34]. However, studies on macrophage-derived EVs in the context of disease and repair have predominantly relied on murine cells or immortalized human macrophage lines, limiting their translational value [19, 35–37]. These systems, while informative, may not fully recapitulate the impact of metabolic modulation. In this study, we addressed this gap, by combining metabolic programming trough PKM2 activation with DASA-58 for the first comprehensive characterization of EVs derived from primary human monocyte–derived macrophages, testing their effects on endothelial cells and mesenchymal stem cells-two cell types that play central roles in angiogenesis and osteogenesis.

Our results demonstrated that polarized human macrophage-derived EVs have distinct biological functions. EVs^M2^ promoted angiogenesis by enhancing endothelial cell migration, proliferation, and tube formation, whereas EVs^M1^ favored osteogenesis by stimulating MSC migration and early ALP activity. These polarization-dependent roles support the idea that macrophage subsets contribute to different phases of tissue repair. However, our findings also diverged from those reported in murine models [35–37]. While murine studies suggested that M2-derived EVs are more pro-osteogenic [22], we found that EVs^M1^ exerted stronger osteogenic effects in human cells. This discrepancy highlights important species-specific differences and stresses the need to validate therapeutic strategies in human-relevant in-vitro systems. Moreover, our data showed that EVs from a single macrophage state lack the regenerative profile needed to promote both angiogenesis and osteogenesis, supporting the development of metabolically engineered EVs with dual regenerative potential.

The analysis of EV miRNA cargo provided mechanistic insights into the observed functional differences. The aim of this approach was to identify miRNAs regulated by macrophage polarization stimuli and to characterize the main miRNAs incorporated into the EV cargo of polarized macrophages. We profiled the expression of EV miRNAs differentially expressed between polarization states. EVs^M1^ were enriched in miRNAs associated with endochondral ossification, consistent with their stimulation of osteogenic processes [38, 39]. In contrast, EVs^M2^ contained miRNAs targeting genes linked to angiogenesis and cell adhesion [40, 41]. Interestingly, only a single miRNA was differentially expressed in EVs^Mφ^ compared with EVs^M1^ and EVs^M2^, and this miRNA has previously been identified as a regulator of monocyte-to-macrophage differentiation [42]. The relatively small number of differences between EVs^Mφ^ and EVs^M2^, compared to previous reports, may represent a limitation of our study [43]. Specifically, the short polarization period used here (4 h) may have reduced the number of detectable changes in miRNA expression, and a longer polarization time could potentially reveal a broader range of differentially expressed miRNAs.

A major novelty of our study lies in the exploration of metabolic modulation as a strategy to refine EV function. PKM2 has emerged as a central regulator of macrophage metabolic reprogramming [44]. Beyond its role in immunometabolism, PKM2 influences mRNA expression, cytokine production, and cellular activation, particularly in response to inflammatory stimuli such as LPS [45, 46]. Of particular relevance, the PKM2 activator DASA-58 was previously shown to induce M2-like mRNA expression in murine macrophages [18]. However, its impact on human macrophages, and especially on their EVs, had not been studied. In this context, we tested whether activation of PKM2 with DASA-58 could generate an EV population that integrates features of both EVs^M1^ and EVs^M2^. Our findings showed that DASA-58 shifted macrophage polarization markers: pro-inflammatory mediators such as *CXCL10* and *CD38* decreased, while *IL10* expression increased. A similar effect had previously been reported for the PKM2 activator TEPP-46 in murine macrophages [46]. At the metabolic level, DASA-58 reduced oxidative phosphorylation and basal respiration, while also downregulating glycolytic mRNAs including *PKM2*, *HK2*, and *HIF1*_α_ [18]. These changes produced a hybrid macrophage phenotype, which was reflected in a distinct EV profile.

Functionally, EVs from DASA-58–treated macrophages (EVs^100D^) promoted endothelial migration and tube formation, effectively mimicking the pro-angiogenic activity of EVs^M2^. By contrast, they did not alter osteogenic activity, retaining the early pro-osteogenic properties seen with EVs^M1^. EVs^100D^ displayed a unique miRNA signature, characterized by reduced levels of pro-inflammatory miRNAs typically enriched in EVs^M1^, while simultaneously incorporating a pro-angiogenic EV miRNA cargo, hsa-miR-423-3p, which we also observed in EVs^M2^ cargo [40, 47]. Importantly, EVs^100D^ did not simply represent a mixture of M1- and M2-derived cargo; instead, they exhibited a novel profile shaped by metabolic modulation, underscoring the potential of targeted interventions to direct EV composition. GO enrichment analysis provided further insight: whereas EVs^M1^ were associated with “regulation of mesenchymal stromal cell differentiation”, EVs^100D^ were enriched for processes linked to “osteoblast differentiation’. This pattern suggests that the hybrid EV profile produced by DASA-58 treatment may help bridge early and late stages of regeneration, reflecting the physiological sequence of endochondral ossification [48]. Such a mechanism may also explain why we did not observe significant changes in late-stage osteogenesis (e.g., calcium deposition) within our experimental timeframe.

Taken together, our study shows that EVs released from M1 and M2 macrophages reflect the complementary roles of their parent cells. EVs^M1^ supported early steps of bone formation, while EVs^M2^ promoted blood vessel growth, though their effects were not identical to what has been reported in mouse models. Importantly, we show that changing macrophage metabolism with DASA-58 can reshape EV cargo and produce hybrid vesicles that combine useful features from both M1- and M2-derived EVs. This ability to adjust EV composition through metabolic control offers a promising way to design EV-based therapies that match the different stages of tissue repair. Future studies should test these engineered EVs in pre-clinical models and explore their potential in other types of tissue healing.

## 4. Conclusion

In conclusion, this study provides the first evidence that human macrophage-derived EVs recapitulate the complementary roles of macrophage subsets in tissue repair. Remarkably, activation of macrophage metabolism with DASA-58 reprogrammed M1 macrophages to release angiogenic EVs without compromising their osteogenic support. These findings highlight EVs as active effectors of macrophage function and underscore macrophage metabolism as a promising lever to generate tailored EV populations. Together, they establish metabolic reprogramming as a novel strategy to engineer human macrophage-derived EVs with enhanced therapeutic potential for tissue regeneration.

## 5. Experimental Methods

### Cell culture of human monocyte derived primary macrophages (hMDMs)

Human monocyte-derived macrophages (hMDMs) were isolated from healthy female donors (aged <40 years) using human buffy coats obtained from the Irish Blood Transfusion Service (research ethics approval from the RCSI, REC number: 202303001). The blood was diluted 1:2 in PBS (Sigma, D8537) and layered onto Lymphoprep (StemCellTech, 7851). After centrifugation at 800 × g for 20 minutes, the peripheral blood mononuclear cell (PBMC) layer was collected, and monocytes were isolated from PBMCs using the Magnisort Human CD14 Positive Selection Kit (Invitrogen, 8802-6834-74). The isolated monocytes were then resuspended in hMDM media, which consisted of RPMI 1640 Medium with GlutaMAX™ Supplement (Gibco, 61870036), 10% human serum (from male AB plasma, Sigma, H4522), and 1% Penicillin/Streptomycin (100 U/mL, Sigma, Cat. P4333). The cells were seeded at a density of 5 × 10 cells per well in a 24-well Corning Costar untreated plate and incubated at 37°C in a 5% CO atmosphere for 7 days. Flow cytometry analysis confirmed that 95% of the cells were CD14+CD11b+ (Supplementary Fig.1).

### RNA Extraction and RT-qPCR

Total RNA was extracted from treated hMDMs using Trizol Reagent (Invitrogen, 15596018) and reverse transcribed (High Capacity cDNA Reverse Transcription Kit, 4368813, Thermo Fisher Scientific) using ProFlex system. A real-time quantitative polymerase chain reaction (RT-qPCR) was performed using ChamQ Universal SYBR qPCR Master Mix (Vazyme, Q711-03) on an Applied Biosystems QuantStudio 7 Pro Real-Time PCR System. The primer sequences used are listed in Supplementary Table 1.

### Cytokine Measurement

Cell supernatants from treated hMDMS were centrifuged at 300g for 10 minutes to remove cell debris. ELISA assays were performed using Invitrogen Human IL-8 (#88-8086-88), IL-6 (#88-7066-88), and IL-10 (#88-7106-88) uncoated ELISA kits, following the manufacturer’s protocols.

### Seahorse Metabolic Flux Assay

The oxygen consumption rate (OCR) of hMDMs were assessed using the Agilent Seahorse XF Cell Mito Stress Test and analyzed on an XFe96 Analyzer. To prepare the cells, hMDMs were detached using Versene (Gibco, 15040066) and seeded into a Seahorse 96-well microplate at a density of 2 × 10 cells per well, allowing them to adhere overnight. The following day, cells were polarized to M1 or M2 phenotypes for 3 hours, after which the medium was replaced with XF RPMI-1640 pH 7.4 medium (Agilent) supplemented with 1 mM sodium pyruvate (Agilent), 2 mM glutamine, and 1 M glucose (Agilent). The plates were then incubated at 37 °C in a CO -free environment for 1 hour. Reagents were prepared and loaded into the designated ports of the utility plate, including oligomycin (1.5 µM) to inhibit ATP synthase (complex V), FCCP (0.5 µM) as an uncoupler that disrupts the proton gradient, rotenone/antimycin A (0.5 µM) as inhibitors of complexes I and III, and 2-deoxy-D-glucose (2-DG, 25 mM). The utility plate was initially run on the flux analyser for calibration. After calibration, it was replaced with the Seahorse 96-well cell culture plate, and the experiment proceeded on the Seahorse XFe96 Analyzer using the Mito Stress test protocol provided in the Agilent Seahorse Analytics XF software.

### Cytoplasmic and Nuclear Fractionation

For the localization of PKM-2, hMDMs treated with the relevant conditions were first detached with Versene solution (Gibco, 15040066). The cytoplasm and nuclei were subsequently separated using the NE-PER™ Nuclear and Cytoplasmic Extraction Reagents (Thermo Scientific, 78833). Protein concentration in both cytoplasmic and nuclear fractions was determined with a BCA Protein Assay Kit (Thermo Fisher Scientific, 23227). 2 µg sample of each extract was mixed with NuPAGE™ LDS Sample Buffer (4X) (Thermo Fisher Scientific, NP0007) and NuPAGE™ Sample Reducing Agent (10X) (Thermo Fisher Scientific, NP0004), followed by heating at 95°C for 5 minutes. Protein separation was performed using NuPAGE™ Bis-Tris Mini Protein Gels (4–12%, 1.0–1.5 mm) (Thermo Fisher Scientific, NP0321BOX), and the proteins were transferred onto Immobilon®-P PVDF membranes (Merck, IPVH00010). After the transfer, the membranes were incubated with antibodies specific to PKM-2 (Cell Signaling, 4053S, 1:1000), β-Tubulin (Cell Signaling, 2146S, 1:1000), and Histone-H3 (Cell Signaling, 4499S, 1:2000). The blots were detected through chemiluminescence imaging using the Amersham imaging system.

### Macrophage polarization and metabolic modulation with PKM-2 allosteric activator DASA-58

hMDMs were polarized into the pro-inflammatory M1 phenotype by treating them with lipopolysaccharide (LPS) from *Escherichia coli* O111 (100 ng/mL) (Sigma, L2630) and interferon-γ (IFN-γ) (20 ng/mL) (PeproTech, 300-02-20UG) for 24 hours. The Mφ control cells remained untreated, while the M2 control cells were polarized to the anti-inflammatory M2a phenotype using interleukin-4 (IL-4) (20 ng/mL) (PeproTech, 200-04-5UG) and interleukin-13 (IL-13) (10 ng/mL) (PeproTech, 200-13-10UG) for 24 hours. For metabolic modulation, hMDMs were treated with the PKM-2 activator DASA-58 (MedChemExpress, HY-19330) at concentrations of 50 µM and 100 µM for 2 hours. This treatment aimed to promote PKM2 tetramerization, after which the cells were polarized to the M1 phenotype with LPS and IFN-γ.

### Collection of conditioned media (CM) and extracellular vesicle (EV) separation

After hMDMs were polarized and metabolically modulated with DASA-58, the cells were washed twice with PBS, and the medium was replaced with fresh, serum-free medium. The following day, conditioned media (CM) were collected and processed by sequential centrifugation: first at 300 × g for 10 minutes to remove cell debris, followed by 2000 × g for 20 minutes to eliminate apoptotic bodies. The CM was then concentrated using Amicon® Ultra-15 centrifugal filter tubes with a 3 kDa molecular weight cutoff (Merck, UFC9003) and subsequently filtered through 0.45 µm filters. EVs were separated from the concentrated CM by differential ultracentrifugation. This was performed using a Sorvall WX 100+ Ultracentrifuge equipped with a swinging bucket rotor (TH600) at 110,000 × g for 90 minutes, with maximum acceleration and deceleration settings. The resulting pellets were washed with PBS and subjected to a second ultracentrifugation cycle under the same conditions. Finally, the EV pellets were resuspended in 50 µL of sterile, filtered PBS. Protein concentration of the EVs was measured using a BCA Protein Assay Kit (Thermo Fisher Scientific, 23227).

### Nanoparticle Tracking Analysis

EV concentration and size were determined using a NanoSight NS300 system (Malvern Technologies) equipped with a 488 nm laser. Samples were diluted in filtered PBS at a ratio of 1:250–1:500 prior to injection. Five readings were performed, each lasting 60 seconds, with the camera level set to 14. Data were analyzed using NTA 3.4 software with a detection threshold of 5.

### Western Blot of EVs

For EV characterization, 10x RIPA buffer (Cell Signaling Technology, 9806S) with Protease/Phosphatase Inhibitor Cocktail (100X) (Cell Signaling Technology, 5872) was added to cells or EV suspensions. The protein concentration of lysed samples was measured using the BCA Protein Assay Kit (Thermo Fisher Scientific, 23227). Two µg of each sample were prepared with NuPAGE™ LDS Sample Buffer (4X) and NuPAGE™ Sample Reducing Agent (10X), then heated at 95°C for 5 minutes. Protein separation was carried out using NuPAGE™ Bis-Tris Mini Protein Gels (4–12%, 1.0–1.5 mm), and transfer was performed onto Immobilon®-P PVDF membranes. Following transfer, the membranes were incubated with the following specific antibodies: anti-syntenin (1:1000 dilution, ab133267, Abcam), anti-Grp94 (1:1000 dilution, ab238126, Abcam, USA), anti-flotillin-1 (1:10,000 dilution, ab133497, Abcam, USA). The blots were then imaged by chemiluminescence using the Amersham imager.

### Transmission Electron Microscopy

EV grids (Formvar/Carbon 300 mesh, Copper approx. grid hole size: 63µm, Ted Pella Inc.) were incubated with EV samples for 20 minutes. Grids with adhered vesicles were rinsed in 0.05 M Phosphate buffer and negatively stained with 1% uranyl acetate. EV imaging was conducted via a JEOL JEM1400 transmission electron microscope (TEM) coupled with an AMT XR80 digital acquisition system [49].

### EV Phenotyping by flow cytometry

EV phenotyping was performed using the MACSPlex Human Exosome Kit (Miltenyi, 130-108-813) following the overnight protocol provided by the manufacturer, utilizing 1.5 mL reagent tubes as specified in the instructions. For these experiments, 2 µg of EVs were used. Flow cytometry was performed using the Aurora flow cytometer (Cytek Biosciences), and the data were analyzed using FlowJo Software version 10.

### Human Umbilical Vein Endothelial Cell (HUVEC) and Human Bone Marrow Mesenchymal Stromal Cell (hMSC) Culture

Human umbilical vein endothelial cells (HUVECs) (Promocell, C-12203) and Green Fluorescent Protein (GFP) -HUVECs (Angio-proteomie, cAP-0001GFP) were cultured in endothelial cell basal medium (EBM) plus endothelial cell growth medium supplements (Promocell, C-22011) supplemented with 10% FBS (Gibco) and 1% Penicillin/Streptomycin (100 U/mL, Sigma, P4333) and incubated at 37°C in a humified 5% CO_2_ atmosphere. HUVECs were used at maximum passage number 8. Primary human MSCs (Lonza) were cultured with Dulbecco’s modified Eagle’s medium (DMEM) GlutaMAX (Gibco) supplemented with 10 % (v/v) FBS (Gibco) and 1 % P/S (Sigma-Aldrich) and used at maximum passage number 4.

### Scratch Assay

For the scratch assay, culture inserts (Ibidi, 80209) were used. HUVECs and MSCs were trypsinized, and a cell suspension of each cell type with a density of 3 × 10 cells/mL was prepared. For each side of the cell culture inserts, 70 µL of the cell suspension was applied. Cells were incubated overnight at 37°C and 5% CO to allow adherence. The following day, the culture inserts were gently removed, and the remaining growth media were carefully aspirated. For HUVEC migration, 300 µL of 1 µg/mL EVs in EBM supplemented with 10% EV-depleted FBS were applied. For MSCs, 300 µL of 1 µg/mL EVs in Serum-Free DMEM-Glutamax were used. A Celldiscoverer 7 microscope (Carl Zeiss Ltd., Cambridge, UK) equipped with a 5x objective and 2x optovar was used to record phase gradient contrast images with an Axiocam 506 camera. Each position was imaged every hour for 48 h. The resultant time-lapse images were analyzed using an Ilastic [50] version 1.4.0 pixel classifier trained to detect the [wound/scratch] area. In FIJI [51] the Ilastik model was run on images using the “Run Pixel Classification Prediction” command that comes with the Ilastik plugin for FIJI. The resultant prediction had a gaussian blur applied and any holes filled before measurement of the final area for all timepoints. All images where prepared in FIJI.

### BrDu Cell Proliferation Assay

HUVECs and MSCs were resuspended in their complete growth media and seeded at 1x 10^4^ cells per well in 96-well plates, followed by overnight incubation to allow for cell adhesion. The next day, cells were synchronized before EV treatments by replacing the complete growth media with serum-free media (EBM for HUVECs, DMEM-Glutamax for MSCs) for 24 hours. On the following day, cells were treated with 100 µL of 1 µg/mL EVs for an additional 24 hours. Cell proliferation was assessed using the Cell Proliferation ELISA, Bromodeoxyuridine (BrdU) assay (Roche, 11647229001) following the manufacturer’s instructions.

### Tube Formation Assay

The tube formation capacity of HUVECs was assessed using a 96-well Angiogenesis μ-slide system (Ibidi, 89646). Each well was pre-coated with 10 µL of Geltrex (ThermoFisher, A1413201) and allowed to polymerize at 37°C for 30 minutes. HUVECs (1 × 10 cells) were then seeded onto the polymerized matrix and incubated at 37°C for 9 hours in the presence of EVs. Images were acquired using a Celldiscoverer 7 microscope (Carl Zeiss Ltd., Cambridge, UK) equipped with a 5× objective, a 2× optovar, and an Axiocam 506 camera. To capture the curvature of the matrix surface, Z stack images composed of 22 slices were used to capture the depth in the sample over a range of 966 µm for each position. Expressed GFP was excited with a 470 nm LED and the emission collected with a quad bandpass filter with the 501-547nm band. All Z stack images were processed with CLIJ2 in FIJI [52]. The ‘Extended Depth of Focus - Variance projection on GPU’ command with a radius_x and radius_y value of 2.0 and a sigma value of 10 was used to yield a single image for subsequent analysis. Tube formation was analyzed using ImageJ software with the Angiogenesis Analyzer plug-in[53].

### ALP Activity and Alizarin Red Staining

hMSCs were seeded at 1.2 × 10 cells per well in a 48-well plate for ALP activity and 2 × 10 cells per well in a 24-well plate for alizarin red staining. After 48 hours, the culture medium was replaced with osteogenic differentiation medium (DMEM + GlutaMAX supplemented with 100 nM dexamethasone, 10 mM β-glycerophosphate, and 0.05 mM L-ascorbic acid-2-phosphate) containing 1 µg/mL EVs. The medium was refreshed on days 4, 7, and 11. ALP activity was quantified on day 7 using the SigmaFast p-nitrophenyl phosphate (pNPP) kit (Sigma Aldrich, N1891) following the manufacturer’s instructions. To normalize ALP activity, total DNA content was measured using the QuantiT™ PicoGreen™ dsDNA Assay (Invitrogen, P7589). To assess mineral deposition, cells were fixed with 4% paraformaldehyde (PFA) for 10 minutes. Following fixation, cells were washed twice with PBS and once with deionized water (diH O). 1 mL of 1% Alizarin Red Stain (ARS) was added to each well and incubated for 20 minutes at room temperature. Excess stain was removed through sequential washes in diH O, and wells were imaged using an Olympus IX83 inverted microscope equipped with a 10× objective with Olympus U-TV0.63XC camera.

### miRNA sequencing and data analysis

Total RNA was extracted from EV suspension using the miRNeasy Mini kit (Qiagen) with the following modifications: EVs were lysed by addition of 1 mL Qiazol containing the mix of miRCURY spike-in controls (Qiagen) which were diluted 1:250 below the recommended concentration to avoid overrepresentation. Isopropanol precipitation was enhanced by addition of glycogen (Thermo Fisher) at a final concentration of 50 µg/ml. The bound RNA was washed twice on a Qiacube liquid handling robot and eluted in 30 µl nuclease-free water. Total RNA was stored at -80°C until further analysis. For small RNA-sequencing, 8.5 µl were mixed with 1 µl miND® spike-in controls (TAmiRNA) and used for library preparation using a single-adapter ligation and circularization kit (RealSeq Biosciences) as described before [54]. Between 17 and 19 PCR cycles were applied for library amplification and subsequently quantified on a Fragment Analyzer using a high sensitivity RNA chip (Agilent). Libraries were pooled at equimolar rate based on the microRNA peak concentration, and size purified on a BluePippin (Sage Science). Libraries were sequenced on an Illumina NextSeq 2000 platform using a P2 flow cell in single-read mode with 100 bp read length. Next-generation sequencing (NGS) data was analyzed using the miND® analysis pipeline [54]. Statistical analysis of preprocessed NGS data was done with R v4.0. Differential expression analysis with edgeR v3.32 used the quasi-likelihood negative binomial generalized log-linear model functions provided by the package [55]. The independent filtering method of DESeq2 was adapted for use with edgeR to remove low abundance miRNAs and thus optimize the false discovery rate (FDR) correction [56]. Additional NGS QC and absolute quantification of miRNAs was done using miND® spike-ins based on a linear regression model. For data visualization, Plotly, Hiplot and ClustVis were used [57–59]. For target prediction and functional enrichment analysis, the tool miRNAtap v1.40.0 was used to derive targets for selected miRNAs. Results from five different miRNA-mRNA interaction databases DIANA, Miranda, PicTar, TargetScan and miRDB are aggregated and targets are filtered by specifying a minimum number of three databases where the target gene must be found. Based on the miRNA targets enriched biological processes (BP) and molecular functions (MF) were calculated using the tool topGO v2.58.0. Gene ontology (GO) annotations were derived from org.Hs.eg.db v3.20.0. Fisher test using the elim algorithm was used to test for enrichment. All GO relevant genes, annotated in the human genome were used as a reference dataset. Interactions between miRNAs and target genes were visualized as Chorddiagrams using the package chorddiag v0.1.3.

### Statistics

Data are presented as mean□±□SD, based on at least three independent donor replicates for each assay, and were analyzed using GraphPad Prism 10 (GraphPad Software, Inc.). Unless otherwise specified, statistical analysis was performed using one-way ANOVA followed by Tukey’s multiple comparisons test. For the MACSPlex analysis, a two-way ANOVA with Tukey’s multiple comparisons test was used. For all analyses, p <□0.05 was considered statistically significant, with the following notation applied: ****p < 0.0001, ***p < 0.001, **p < 0.01, *p < 0.05.

## Acknowledgements

This project is funded by the European Union under Marie Sklodowska-Curie Post-doctoral Fellowship grant agreement No 101106209 (METABOLATE) and by Research Ireland through the Frontiers for the Future Project Grant (19/FFP/6533) and Award (23/FFP-A/12166). The material presented, and views expressed here are the responsibility of the author(s) only. The EU Commission takes no responsibility for any use made of the information set out.

## Ethics Statement

Human buffy coats obtained from the Irish Blood Transfusion Service (research ethics approval from the RCSI, REC number: 202303001).

## Data availability statement

The data are available from the corresponding author(s) on reasonable request.

## Author contributions

**C.G.** conceptualization, validation, formal analysis, investigation, visualization, funding acquisition and writing-original draft, writing-review and editing; **P.K.** validation, formal analysis, investigation, visualization, writing-review and editing; **C.S.M.** validation, formal analysis, investigation, visualization of osteogenesis experiments; **C.P.** validation, formal analysis, investigation, visualization of flow cytometry experiments, writing-review and editing; **B.L.C** formal analysis, investigation, data curation and visualization of imaging experiments, writing-review and editing; **M.P.** formal analysis, data curation and visualization of miRNA sequencing: **M.H.** formal analysis, data curation, resources, supervision, writing-review and editing: **A.M.C.** conceptualization, resources, supervision, funding acquisition and writing-original draft, writing-review and editing; **D.A.H**. conceptualization, resources, supervision, funding acquisition and writing-original draft, writing-review and editing. All authors approved the submission of the manuscript.

