## Supplemental Material for "Metabolic Reprogramming of Human Macrophages Drives the Formation of Hybrid M1/M2 Pro-Regenerative Extracellular Vesicles"

Funding: This project is funded by the European Union under Marie Sklodowska- Curie Post-doctoral Fellowship grant agreement No 101106209 (METABOLATE) and by Research Ireland through the Frontiers for the Future Project Grant (19/FFP/6533) and Award (23/FFP-A/12166).

Keywords: extracellular vesicles; human macrophages; reprogramming; angiogenesis; miRNA; osteogenesis

<u>**SUPPLEMENTARY FIGURES**</u>

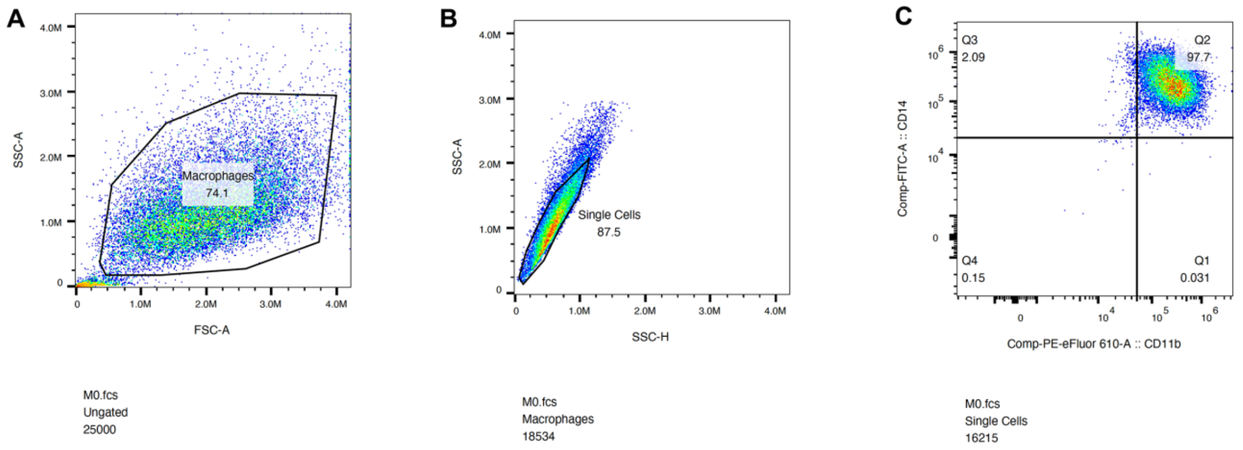


**Supplementary Figure 1: Flow cytometry gating strategy for CD14$^{+}$CD11b$^{+}$ macrophages. (A)** Events were plotted on SSC-A vs. FSC-A to exclude debris and define the cell population. **(B)** Singlets were gated using SSC-A vs. SSC-H. **(C)** Representative plot showing CD14$^{+}$CD11b$^{+}$ macrophages within the singlet gate.

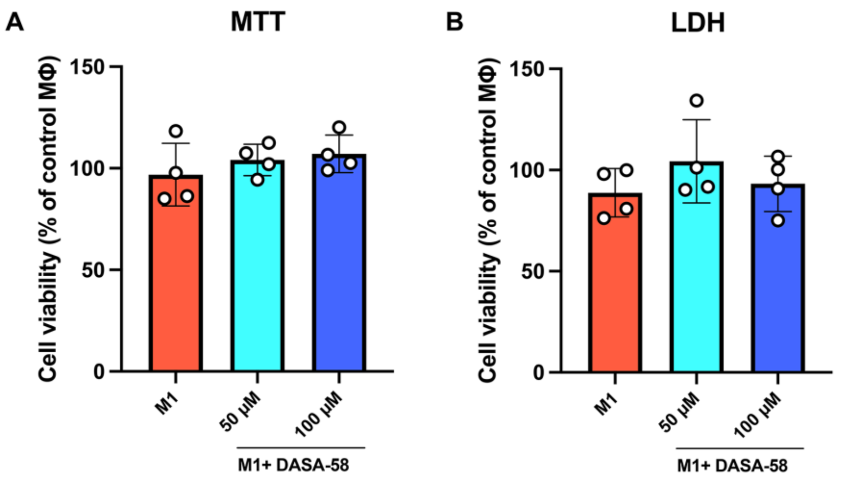


**Supplementary Figure 2: Preliminary cytotoxicity assay confirmed that both concentrations of DASA-58 were well tolerated by the human macrophages, with no detectable cytotoxic effects. (A)** MTT assay and LDH assay **(B)** in human macrophages after pre-treated with DASA-58 for 2h and then activated with LPS+IFNγ for 22h. Error bars in the graphs represent mean ± SD of four independent donors.

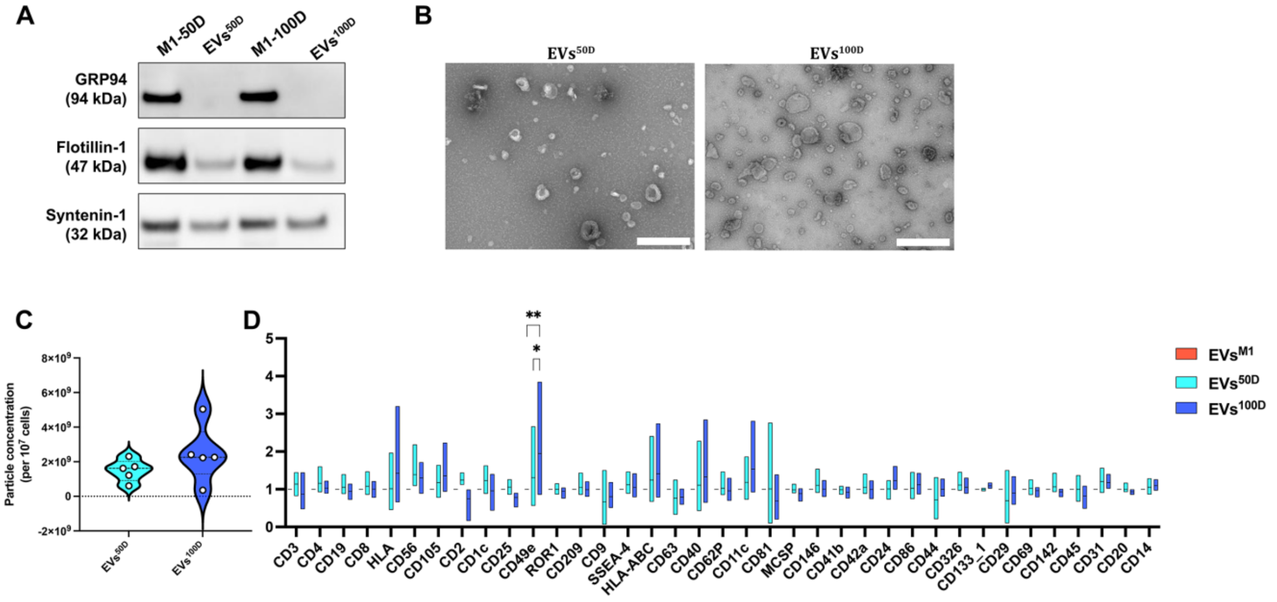


**Supplementary Figure 3: EV characterization of EVs$^{50D}$ and EVs$^{100D}$.** **(A)** Representative Western-blot analysis of non-EV marker GRP94 and, EV markers Flotilin-1 and Syntenin-1. **(B)** Representative transmission electron microscopy images of macrophage EVs. Scale bar=500 nm. **(C)** Quantitative analysis of EV concentration by Nanoparticle tracking analysis. N=5 donors. **(D)** Surface marker profiling of EVs with MACSplex Exosome Kit. **p < 0.01, *p <0.05 (two-way ANOVA and Tukey multiple comparison). N = 4 donors.

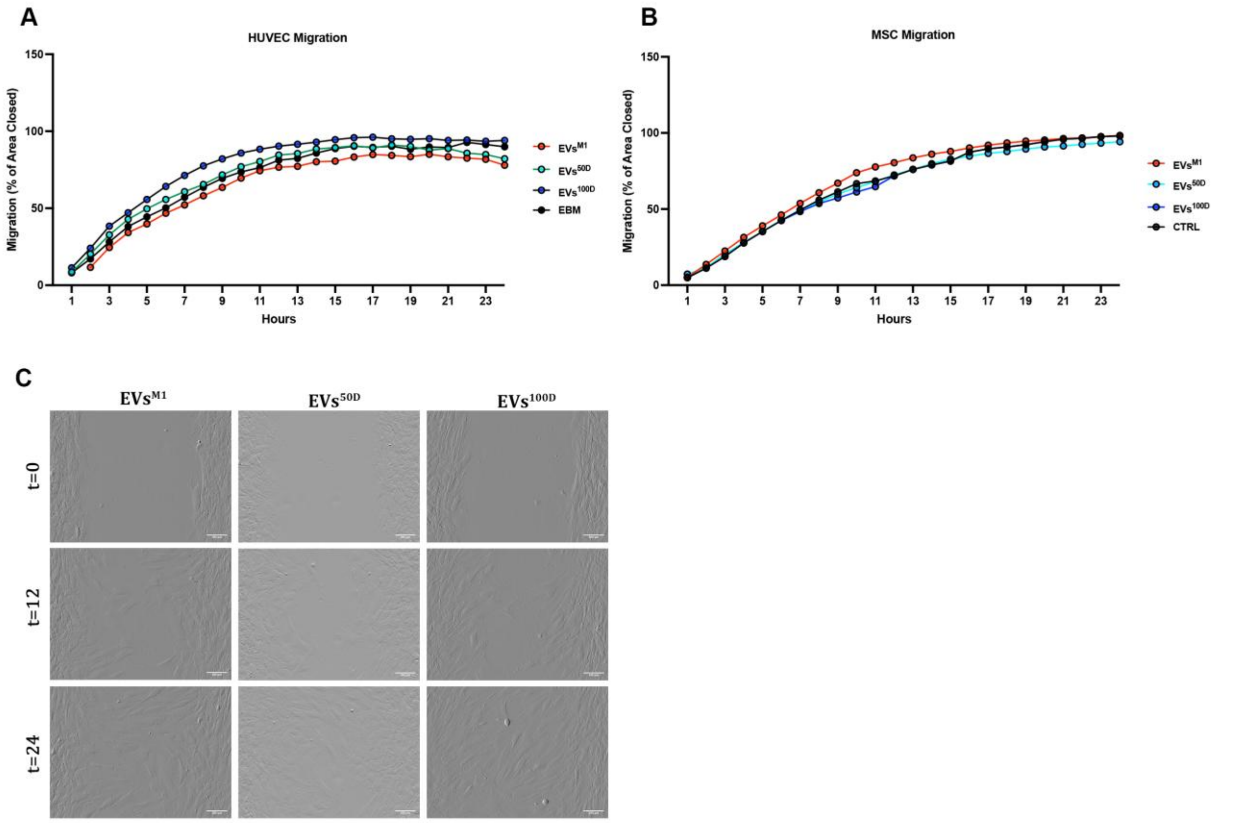


**Supplementary Figure 4: (A)** Plot graph of MSC migration and **(B)** HUVEC migration for 24 hours. **(C)** Representative images of MSC migration. Scale bar=100 µm.

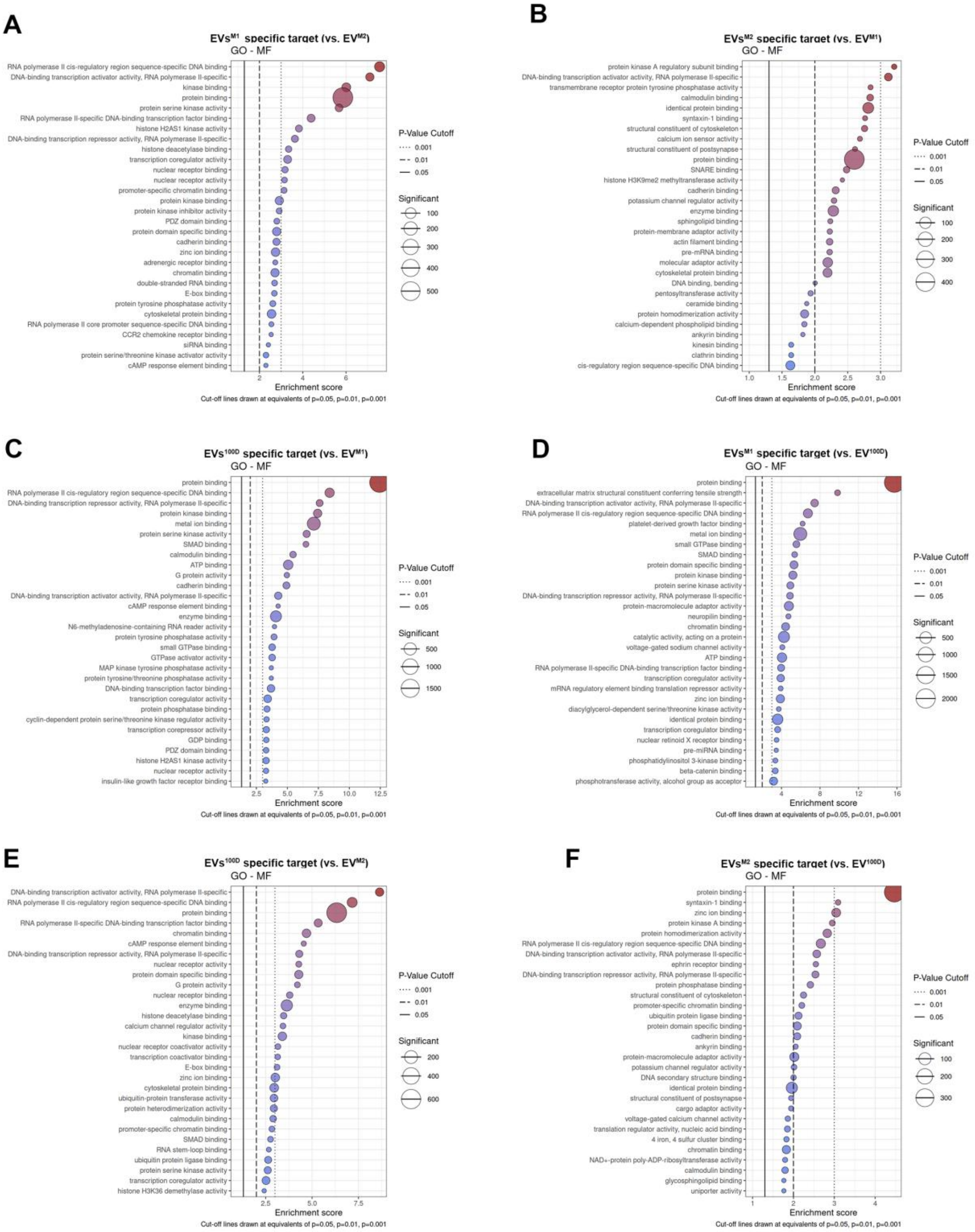


**Supplementary Figure 5: Top 30 significantly enriched GO molecular functions associated with all differentially expressed miRNAs**. **(A)** $EVs^{M1}$ vs $EVs^{M2}$ ($EVs^{M1}$ specific miRNA subset) , **(B)** $EVs^{M2}$ vs $EVs^{M1}$ ($EVs^{M2}$ specific miRNA subset), **(C)** $EVs^{100D}$ vs $EVs^{M1}$ ($EVs^{100D}$ specific miRNA subset), **(D)** $EVs^{M1}$ vs $EVs^{100D}$($EVs^{M2}$ specific miRNA subset), **(E)** $EVs^{100D}$ vs $EVs^{M2}$ ($EVs^{100D}$ specific miRNA subset), **(F)** $EVs^{M2}$ vs $EVs^{100D}$ ($EVs^{M2}$ specific miRNA subset). N=5 donors.

### SUPPLEMENTARY TABLE

**Table S1: Forward and reverse SYBR Green primer sequences for quantitative real-time PCR.**

| Gene name | Forward/Reverse sequence |
|---|---|
| **18S** | F: 5’- GGCGCCCCCTCGATGCTCTTAG<br>R: 5’- GCTCGGGCCTGCTTTGAACACTCT |
| **CXCL10** | F: 5’- GGTGAGAAGAGATGTCTGAATCC<br>R: 5’- GTCCATCCTTGGAAGCACTGCA |
| **CD38** | F: 5’- TCTTGCCCAGACTGGAGAAAGG<br>R: 5’- TGGACCACATCACAGGCAGCTT |
| **VEGF-A** | F: 5’- AGGGCAGAATCATCACGAAGT<br>R: 5’- AGGGTCTCGATTGGATGGCA |
| **CD206** | F: 5’- CTACAAGGGATCGGGTTTATGGA<br>R: 5’- TTGGCATTGCCTAGTAGCGTA |
| **IL-10** | F: 5’- AAGCCTGACCACGCTTTCTA<br>R: 5’- ATGAAGTGGTTGGGGAATGA |
| **PKM2** | F: 5’- ATGGCTGACACATTCCTGGAGC<br>R: 5’- CCTTCAACGTCTCCACTGATCG |
| **HK2** | F: 5’- GAGTTTGACCTGGATGTGGTTGC<br>R: 5’-CCTCCATGTAGCAGGCATTGCT |
| **HIF1alpha** | F: 5’- TATGAGCCAGAAGAACTTTTAGGC<br>R: 5’- CACCTCTTTTGGCAAGCATCCTG |
